# Selective packaging of pathogenic non-coding RNAs into plant exosomes mediated through Phloem Protein 2 and Tetraspanin 8 may represent a key mechanism for their cross-kingdom transmission

**DOI:** 10.1101/2025.03.19.644080

**Authors:** Neha Devi, Nisha Devi, Vasudha Sharma, Ravi Gupta, Sunny Dhir, Yashika Walia

**Affiliations:** Molecular Plant Virology Group, R&D Lab, Department of Bio-Sciences & Technology, Maharishi Markandeshwar (Deemed to be University), Mullana, Ambala-133207 HR India; Plant Stress Physiology and Proteomics Laboratory, College of General Education, Kookmin University, Seoul, South Korea

**Author notes:** Corresponding Authors, Dr. Yashika Walia;, Dr. Sunny Dhir.

**Keywords:** Extracellular vesicles, Viroid RNA, Phloem Protein 2, Tetraspanin 8, Exomotifs, m6A methylation

## Abstract

The plant apoplast and extracellular vesicles (EVs) contain diverse RNA species, including small RNAs, long non-coding RNAs, and circular RNAs, many of which cross kingdom boundaries through EV-mediated transport. Viroid RNAs—pathogenic circular non-coding RNAs—are known to move from plants to fungi and insects, though the mechanism remains unclear. Using apple scar skin viroid (ASSVd) and cucumber as a host, we demonstrate that viroid RNAs in circular, linear, and small RNA forms are present in cucumber EVs. ASSVd RNA exhibits features of extracellular RNAs, such as m6A methylation and five EXOmotifs. Through mass spectrometry and immunoblotting, we identify tetraspanin 8 (CsTET8) as a conserved EV marker and report CsPP2, a phloem lectin with an RNA-binding domain, as a novel EV-associated protein. RNA co-immunoprecipitation and immuno-enrichment assays reveal that CsPP2 and ASSVd co-localize in CsTET8-positive EVs, with direct interactions between CsPP2-CsTET8 and CsPP2-ASSVd. We propose CsPP2 functions as an RNA-sorting protein that mediates RNA loading into EVs. Further, we show that whiteflies can acquire ASSVd via EVs and transmit it to healthy plants, indicating a cross-kingdom transmission route. These findings establish viroid RNAs as a model system to study the molecular basis of RNA trafficking across biological kingdoms.

## Introduction

The role of the apoplast and EVs in mediating cross-kingdom RNA trafficking has gained attention recently (Zhang et al., 2016, Wang et al., 2024). EVs are small, membrane-bound vesicles secreted by cells that can encapsulate a variety of biomolecules, including RNA (Zhang et al., 2016; He et al., 2021, Karimi et al., 2022; Wang et al., 2024). Plants are known to contain three classes of EVs (Ambrosonethe et al., 2023). The first class is represented by exosomes with sizes ranging from 30-150nm which are specifically enriched in tetraspanin 8 (TET8), an ortholog of mammalian CD63, CD81, and CD151 (Chen et al., 2022, Jankovičová et al., 2020). The second class is represented by micro-vesicles with sizes ranging from 150-300nm and are associated with syntaxins proteins, of which Penetration 1 (PEN1) is well characterized (Rutter et al., 2017). The third class is represented by EXPO vesicles which are double membrane bodies fused with the plasma membrane (PM), releasing a single membrane vesicle into the cell wall (Wang et al., 2010). Exo70E2 is essential for exocyst subunit recruitment and EXPO formation in both plants and animals (Ding et al., 2014).

Plant apoplast and EVs are enriched in various RNA species, such as long non-coding RNAs (lncRNA), circular RNAs, (circRNA) micro RNAs (miRNAs), and tasiRNAs (Karimi et al., 2022). The process of cross-kingdom RNA transfer via EVs was first observed during the interaction between Arabidopsis and *Botrytis cinerea*, where pathogen-derived small RNAs (sRNAs) were shown to travel across kingdoms and interact with plant argonaute (AGO) to target mitogen-activated protein kinase transcripts, promoting pathogen colonization (Weiberg et al., 2013). Subsequent studies revealed that plants can send their small RNAs in TET8-specific exosomes to suppress fungal virulence genes (Cai et al., 2018). In addition, they also carry mRNAs that are translated into fungi and reduce their pathogenicity (Wang et al., 2024). Recent studies have identified various mechanisms involved in the loading of RNA into EVs. Specifically, RNA-binding proteins such as AGO1, annexins, and RNA helicases play significant roles in the incorporation of plant small RNAs into TET8-specific exosomes (He et al., 2021; Karimi et al., 2022). Furthermore, glycine-rich binding protein (GRP7) and AGO2 have been linked to the secretion of N6-methyladenosine (m6A) modified circular and long non-coding RNAs, as well as small RNAs in the apoplast (Karimi et al., 2022). Similar to the animals, circular RNAs in the plant apoplast are notably enriched with m6A modifications (Sabrina Garbo et al., 2024; Karimi et al., 2022). Certain factors have been identified in animals and plants that could assist the sorting of RNAs inside the vesicles. These factors encompass characteristics such as a high GC content, the presence of an EXOmotif (a motif known to facilitate preferential sorting of RNA into exosomes or EVs), and their ability to modify RNAs and interact with RNA binding proteins (O’Grady et al., 2022, Ruben Garcia-Martin et al., 2022, Sabrina Garbo et al., 2024, Xiao-Man Liu 2025).

A class of non-coding circular RNAs is represented by viroids which are RNA-only pathogens infecting plants with distinct structural and functional properties (Flores et al., 2016; Navarro et al., 2021; Hammond, 2017). Despite their small size (246-401nts) and non-coding properties, they autonomously replicate within infected plant cells and cause a variety of diseases. The ability of viroids to infect plants is enabled by their secondary structure (Branch et al., 1984; Daros et al., 1994; Warrilow et al., 1999; Navarro et al., 2000; Rodio et al., 2007), which is crucial for their replication and movement and allows hijacking of plant cellular machinery for their propagation (Schnölzer et al., 1985; Schmitz and Riesner, 1998; Sano et al., 1992; Reanwarakorn & Semancik, 1998). Recent studies have shown that viroids are not limited to infecting plants but can also affect other biological kingdoms, including fungi, insects, and bacteria (Tian et al., 2020; Walia et al., 2015; Leichtfried et al., 2022; Afanasenko et al., 2022). This cross-kingdom infection potential suggests that viroids, much like other RNA entities, can traverse biological barriers and may play a role in complex ecological interactions (Cai et al., 2019, Wang et al., 2024). For example, research on apple scar skin viroid (ASSVd) demonstrated that viroids get transmitted from infected apple trees to fungi, such as *Alternaria alternata*, a common plant pathogen (Tian et al., 2022). We have previously reported that the glass-house whitefly, *Trialeurodes vaporariorum*, mediates the transmission of ASSVd and cucumber PP2 assists in its acquisition and transmission (Walia et al., 2015). Moreover, certain aphid species and codling moth larvae, which feed directly on infected apple trees, were found to transmit apple chlorotic fruit spot viroid (ACFSVd), highlighting the role of insect vectors in cross-kingdom viroid transmission (Leichtfried et al., 2022). Potato spindle tuber viroid (PSTVd) transmission from infected plants to *Phytophthora infestans* also emphasizes the inter-kingdom dynamics of viroid spread (Afanasenko et al., 2022).

All these findings together suggest that viroids, as circular non-coding RNAs, may take advantage of RNA sorting and encapsulation mechanisms within EVs to facilitate their intercellular movement and potential cross-kingdom dissemination. By associating with EVs, viroids could bypass the hostile intracellular environment of the host, enabling long-distance transport or transmission across different organisms. Using ASSVd as a model for circular non-coding RNAs and cucumber as its host, we demonstrate that viroid RNAs in various forms are present in cucumber EVs. ASSVd RNA exhibits extracellular RNA signatures, such as m6A methylation and EXOmotifs, which may facilitate EV packaging. Protein identification by mass spectrometry (MS) identifies TET8 as a conserved EV marker and PP2, a phloem lectin with an RNA-binding domain, as a novel E-associated protein. We demonstrate the interaction between ASSVd-PP2-TET8 and speculate that PP2 might assist in RNA loading inside EVs. Lastly, we show that whiteflies can acquire and transmit viroid RNAs via EVs, highlighting a potential cross-kingdom RNA trafficking mechanism.

## Material and Methods

### Plant Growth

*Cucumis sativus* (Var. Summer Green) and *Nicotiana benthamiana* seeds were cultivated in pots containing a mixture of cocopeat, compost, and vermiculite/Perlite in a 1:1:1 ratio. The plants were kept in an insect-proof plant growth chamber equipped with fluorescent tubes under controlled conditions with a temperature of 24±2°C, relative humidity of 70-80%, and a 12-hour light/12-hour dark cycle.

### *In vitro* synthesis of dimeric ASSVd (+) RNA and transcript inoculation

To generate positive-sense RNA transcripts of ASSVd, recombinant plasmids containing the dimeric ASSVd insert in the correct positive orientation within the pGEMT-easy vector were linearized using *Spe*I. In vitro transcription was carried out using the T7 Highscribe Kit (NEB, USA) according to the manufacturer’s protocol. Approximately 100 ng of the RNA transcripts were subsequently used to infect the first true leaf of cucumber plants through mechanical inoculation using carborundum-dust. The water-inoculated plants were used as controls in all experiments.

### RNA extraction and RT-PCR for ASSVd detection

Total RNA was extracted from 100 mg of cucumber leaves using the RNAiso plus (TakaraBio, USA; Walia et al., 2020). Briefly, 100mg of the cucumber leaves (viroid infected and mock) were frozen in liquid nitrogen, ground in Trizol buffer, and incubated with chloroform. After centrifugation, the supernatant was mixed with isopropanol for RNA precipitation and stored at - 20°C. The RNA pellet was then washed with 75% ethanol and eluted in 20 µL of nuclease-free water. The RNA was quantified using Quantus™ Fluorometer (Promega) and 1µg of RNA was used for RT-PCR with viroid-specific primers ASSVdICc and ASSVdICh (Table S1) (Walia et al., 2014). Reverse transcription was performed using verso cDNA synthesis kit (Thermo Scientific, USA) at 42°C for 60 minutes, followed by PCR using Phusion DNA polymerase (Thermo Scientific, USA) under specified cycling conditions for ASSVd detection (Walia et al., 2014). The PCR products were analyzed on 1.5% agarose gel electrophoresis.

### Isolation of AWF

Apoplastic washing fluid (AWF) from cucumber leaves was isolated as described previously (O’Leary et al., 2014). The leaf petioles were excised, followed by washing and drying the leaves (Fig. S1). The leaves were placed in a 60 mL syringe filled with an infiltration buffer (20mM MES, 2mM CaCl_2_, 0.1M NaCl, pH 6.0). Carefully without damaging the leaves, a negative pressure was created by pulling the plunger gently until the leaves turned dark green. The leaves were removed and dried and 3-4 leaves rolled were placed in a 12 mL syringe inside a 50 mL falcon tube. The tube was wrapped with parafilm and centrifuged at 4,000 rpm for 30 minutes to collect the AWF.

### Trypan blue staining of leaves after AWF isolation for cell death analysis

The integrity of the isolated AWF was confirmed by cell death assay by staining the leaves in trypan blue solution using the protocol as described (http://resources.rothamsted.ac.uk).

### RNA isolation and RT-PCR detection of ASSVd from AWF

One ml of isolated AWF was concentrated to 100µl using an Amicon® Ultra Centrifugal Filter, 10 kDa MWCO (Millipore, USA). RNA was isolated using RNAiso plus (Takara Bio, USA). Briefly, 1ml of RNAiso plus was added to 100µl of AWF followed by the addition of the chloroform. The RNA was precipitated using isopropanol and suspended in 30µl of RNase-free water. One microgram of RNA was used for RT-PCR as mentioned earlier.

### Isolation of EVs

Pure-EVs SEC columns were used to isolate EVs from cucumber plants (mock/viroid infected). For EV isolation, 20 mL of isolated AWF was concentrated to 2 mL using an Amicon filter unit (10kDa cutoff) (Millipore, USA) and was subjected to Pure-EVs SEC columns (Hansa Biomed Life Sciences, Estonia) according to the manufacturer’s instructions. Briefly, the column was washed with 3 volumes of 1× PBS buffer to remove preservative buffer residues. The sample containing EVs (up to 2 mL) was then loaded into the column and eluted with 1× PBS buffer as the mobile phase. Fractions collected were labeled 1-7 as void volume, 8-13 as the EV fraction, and 14-23 as the protein sample (Fig. S2).

### Dynamic Light Scattering (DLS) analyses of EVs

The DLS analyses were performed in a Litesizer 500 (Anton Paar, Germany). 10 µL of isolated EVs were suspended in 990 µL of 1× sterile PBS buffer, loaded into a disposable cuvette, and placed in the litesizer 500. For reproducibility, and standardization, the parameter values were fixed as follows; Temp-25℃, equilibration time-30s, processed runs-5, and Time for each run 10s. The size or homogeneity of exosomes was recognized through the radius or the % Intensity, respectively.

### Transmission electron microscopy of EVs

Purified EVs (10 µl) were diluted with an equal volume of 4% paraformaldehyde at 4°C for 30 min, and then carefully placed on a carbon-coated 300-mesh copper grid for 20 min, followed by fixed by 1% glutaraldehyde for 5 min. After two washings, the grids were contrasted with 2% uranyl acetate and then washed again two times. The morphology of isolated exosomes was visualized with TEM (MAKE: JEOL MODEL: JEM 2100 plus).

### Trypsin, RNase treatment, RNA isolation, and RT-PCR from EVs

To determine whether the viroid RNA was present inside or outside EVs, RNase protection assays were performed. Briefly, 100µl of Purified EVs were treated with 25 µg/µL trypsin (Sigma Life Sciences, USA). Samples were incubated at 37°C for 16-18 hours, followed by the addition of 1U of Proteinase K (Invitrogen, USA)/1mM of PMSF (Sigma, USA) to inactivate trypsin. For isolating the RNA from the EV surface the above EV samples were precipitated with 3 volume ethanol and 1/10 sodium acetate overnight followed by isolation of RNA with RNAiso plus next day. For isolating the RNA inside EVs the purified EVs (100µL) were treated with trypsin + trypsin inhibitor + RNaseA + RNase inhibitor to degrade the outside RNA followed by treatment with 1% (v/v) Triton X-100 (Himedia) to rupture the EVs. RNA isolation using RNAiso plus was carried out to isolate the RNA inside EVs. The isolated RNA was quantified and 200ng of the total RNA was used for RT-PCR from the detection of ASSVd from EV^O^ and EV^I^ as described above. The concentration of trypsin and RNase used in the assays was standardized by taking different dilutions of trypsin and RNase for protein degradation and RNA degradation, respectively (Fig. S3 and S4).

### Urea PAGE and Northern Blot for ASSVd detection

Viroid-infected samples from total leaf, AWF, and EVs (EV^O^ and EV^I^) along with mock RNA from leaf were analyzed using Denaturing Polyacrylamide gel electrophoresis (15% polyacrylamide and 7M Urea). The RNA samples (3µg for AWF, EV^O^, EV^I^ 10ug, and 3ug for CL from infected and 3ųg from Mock) were denatured at 65°C in 2x RNA dye (Thermo Scientific, USA) for 10 minutes and were separated in 0.5× TBE buffer for 2h at 90V. The total RNAs were visualized by staining with Diamond Nucleic acid stain (Promega) and then transferred onto a nylon membrane using 1× TBE transfer buffer. The RNAs were cross-linked on a nylon membrane in a UV crosslinker followed by pre-hybridization at 65°C for 3hrs. The membrane was incubated overnight at 65°C in hybridization buffer with gentle shaking along with a 3’ labeled biotin probe. The membrane was rinsed with washing buffer (1× SSC. 0.1% SDS) thrice for 10 minutes at room temperature and blocked in a blocking buffer (0.1% Malic acid buffer (Roche). The membrane was incubated with HRP-conjugated anti-streptavidin antibody (Sigma, USA) at 1:5000 dilutions for 1h. After a final wash in maleic acid and 0.3% Tween-20 containing buffer, the blot was visualized using ChemiDoc (Protein Simple, Biotechne, USA).

### Generation of negative sense 3’-biotin labeled ASSVd probe

A 3’-biotin-labeled RNA probe was prepared and used for viroid detection using northern hybridization. For probe synthesis, a PCR was carried out using ASSVd plasmid as a template and primers FP1 and RP4 to generate negative sense viroid amplicon along with the T7 promoter sequence. The desired amplicon of 330bp was cut and eluted. Two micrograms of gel eluted product was directly used to generate RNA transcript using a T7 in vitro RNA transcription kit (Thermo Scientific, USA). The transcript at the 3’ end was ligated to pcp-biotin (Jena Biosciences) using T4 RNA Ligase (Promega, USA). The reaction was kept at 16°C overnight. The labeled transcript was purified using a Monarch RNA cleanup column (New England Biolabs, USA) and was used for hybridization at a concentration of 100ng/10ml of hybridization buffer (Fig. S5).

### Immunoprecipitation with m6A antibody

Total RNA was isolated from apoplastic fluid, and EVs using RNAisol plus as described earlier and quantified using a Quantus fluorimeter (Promega, USA). For AWF and EVs, 5µg RNA was incubated with an m6A antibody (Epigenetek-A-1801-020) in IP buffer (50mM Tris HCl pH 7.5, 150mM NaCl, 0.5% NP-40) at 4°C for 4-5 hours, followed by the addition of Protein A/G agarose beads (Merck, USA) and incubated overnight. The samples were washed with IP buffer and eluted from beads using RNAiso plus. The isolated RNA was suspended in 100µl of RNase-free water. For spotting on the membrane, 10µl of the RNA was denatured at 65°C for 10 minutes and applied to a nylon membrane (for detecting with ASSVd probe) and cross-linked. The membrane was pre-hybridized, incubated overnight with a negative sense ASSVd-specific biotin probe, washed, and blocked. After incubation with HRP-labelled streptavidin (Sigma, USA), m6A-modified RNA was visualized using a ChemiDoc system (Protein Simple, Biotechne, USA). For detecting with anti m6A antibody the RNA was spotted on the nitrocellulose membrane and the membrane was blocked in 5% skim milk solution with overnight incubation with 1:5000 dilution of anti m6A antibody (Epigenetek-A-1801-020). The next day membranes were washed and incubated with goat anti-rabbit HRP-conjugated secondary antibodies, washed and visualized using ECL substrate (Biorad, USA), and imaged with a ChemiDoc system (Protein Simple, Biotechne, USA) or X-ray film.

### Mass spectrometry

EVs fraction (100µl) was treated with trypsin followed by incubation with trypsin inhibitor and were then treated with 0.1% Triton X-100. The proteins were TCA precipitated and partially resolved by polyacrylamide gel electrophoresis till they entered the resolving gel. The region was excised and sequenced using HR-LCMS, SAIF facility at IIT-BOMBAY. Briefly, the gel slices were subjected to in-gel trypsin digestion followed by desalting and subjected to LC/MS/MS analysis in Q-Exactive Plus Orbitrap high-resolution MS (Thermo Scientific). The MS spectra were acquired at a resolution of 70,000 in a mass range of 350-2500 m/z. Eluted samples were used for MS/MS events (resolution of 17,500), measured in a data-dependent mode for the 15 most abundant peaks (Top15 method).

### Protein identification and GO enrichment tools

The raw MS/MS spectra were analyzed using the software Thermo Proteome Discoverer 2.2 against the Arabidopsis (TAIR) and cucumber database (Cucurbit genomics-http://cucurbitgenomics.org/). The ShinyGO v0.82 (ShinyGO 0.82 (sdstate.edu)) and string tools were used (STRING: functional protein association networks (string-db.org) for GO analyses. Venn diagram was constructed using the software Bioinformatics & Evolutionary Genomics (Draw Venn Diagram (ugent.be).

### Immunoblots

For immunoblots, cell lysate was prepared by grinding 50mg of leaf tissue in 500ųl of protein extraction buffer (50mM Tris HCl pH 7.5, 150mM NaCl, 10mM MgCl_2_, 2.5mM EDTA, 1mM DTT, 10% Glycerol, 1x protease inhibitor, 1mM PMSF). 20ųl of cell lysate, concentrated AWF, and EVs (EVO and EVI) were mixed with 3 µL of 6× SDS loading buffer, heated at 99°C for 10 minutes, and loaded onto a 15% PAGE, and after run proteins were transferred to a nitrocellulose membrane (for dot blots, the dots were applied on the nitrocellulose membrane directly without the SDS dye). Membranes were blocked with 5% skim milk in TBST and incubated overnight at 4°C with primary antibodies (1:1000 for TET8 (PhytoAb-PHY2750A), 1:10000 for CsPP2 (Walia et al., 2015, 1:10000 anti-GFP (BioBharati Life Science, India) and 1:3000 anti-HA (BioBharati, Life Science, India), anti-actin (Sigma) mouse monoclonal 1:20000 dilution. Membranes were washed and incubated with 1:10000 goat anti-rabbit/anti-mouse HRP-conjugated secondary antibodies (BioBharati, Life Science, India). After a final wash, proteins were visualized using the ECL substrate (Biorad, USA) and imaged with a ChemiDoc system (Protein Simple, Biotechne, USA) or X-ray film.

### RNA immunoprecipitation assay (RIP assay)

For RNA immunoprecipitation from the leaf, 1g/ml of viroid-infected leaf tissue was ground in protein extraction buffer supplemented with RNase inhibitor, and 500µl of lysate was used for RNA immunoprecipitation assay. For RNA immunoprecipitation from infected AWF, 5ml of AWF isolated from viroid-infected leaves was concentrated to 500µl solution using an Amicon filter. Both the lysate and AWF were incubated with 5mg of CsPP2 antibody for 4h at 4℃ with rotation followed by the addition of protein A/G agarose beads overnight. The next day the beads were pelleted with centrifugation at 3000 rpm for 5 minutes and washed three times with protein extraction buffer supplemented with RNase inhibitor. The beads were eluted in 100ul of RNAse-free water. Half of the beads were heated at 100℃ for 10 minutes to release bead-bound proteins for subsequent immunoblot. The rest of the beads were proceeded for RNA isolation using RNAiso plus. The RNA was quantified and used for the detection of ASSVd by RT-PCR or loaded on 15% denaturing PAGE to visualize the total immunoprecipitated RNAs using Diamond Nucleic acid stain (Promega) as described earlier.

### Immunoenrichment of TET8 exosomes

The isolated intact EVs were treated with trypsin overnight, followed by the addition of a trypsin inhibitor. Next, RNase was added for one hour, and then RNase inhibitor was used to inactivate the RNase. The intact EVs were incubated with anti-AtTET8 antibody in IP buffer without NP-40 for 4 hours at 4°C with rotation, followed by the addition of protein A/G agarose beads overnight. The next day, the beads were pelleted and washed with IP buffer containing 1%Triton X-100 and RNase inhibitor. The beads were then eluted in RNase-free water, with part of the eluate used for RNA isolation as previously described. The isolated RNA was tested for the presence of ASSVd by RT-PCR, as described earlier. The remaining beads were heated and loaded onto an SDS-PAGE gel for immunoblotting with anti-AtTET8 and anti-CsPP2 antibodies.

### Co-Immunoprecipitation assay

For co-immunoprecipitation assays involving GFP-TET8 and HA-PP2, Nicotiana benthamiana leaves were harvested 48 hours post agroinfiltration. Tissue extracts were homogenized in Protein extraction buffer (PEB) [5 mM Tris-HCl (pH 7.5), 150 mM NaCl, 10 mM MgCl₂, 1 mM EDTA, 1% NP-40, and 1X protease inhibitor cocktail (Sigma-Aldrich, USA)]. After centrifugation to remove cell debris, the supernatant was pre-cleared using IgG agarose beads (Biobharti Life Sciences, India) for 1 hour at 4°C with rotation. The mixture was centrifuged again, and the IgG agarose bead pellet, used as a non-specific binding control, was collected. The resulting supernatant was incubated overnight at 4°C with anti-GFP agarose beads (BioBharati Life Sciences, India). Following centrifugation, the beads were washed three times with PEB buffer, resuspended in 1× SDS loading dye, and prepared for immunoblot analysis.

### Subcellular Localization and Bi-molecular Fluorescence Complementation (BiFC) Assays

*Agrobacterium tumefaciens* strain GV3101 harboring CsTET8 fused to GFP in the binary vector pSITE2CA was infiltrated into mature *N. benthamiana* leaves. At 48 hours post-infiltration (hpi), tissue sections from infiltrated areas were examined using a confocal microscope (NIKON, AIR PLUS). Images were captured using both bright-field and YFP filter settings. For BiFC assays, GV3101 strains carrying CsTET8 and CsPP2 in BiFC-compatible binary vectors (nVENUS and cCFP) were co-infiltrated into *N. benthamiana* leaves in the indicated combinations. Leaf sections were observed under a confocal microscope at 48 hpi. Images were captured using both bright-field and YFP filter settings.

### Dynamics of CsTET8 and CsPP2 upon viroid infection

Three cucumber plants were infected with ASSVd transcripts along with mock as control. The samples were harvested at 7dpi, 14dpi, 21dpi and 28dpi. The lysates were prepared in 8M urea, mixed with SDS dye, heated, and loaded on 15% PAGE. The proteins were transferred to the membrane and immunoblotted with CsPP2 and AtTET8 antibodies as described earlier.

### Bioinformatic analysis of the ASSVd-CsPP2-CsTE8 interactions

The amino acid sequences of the proteins were obtained from the following accession numbers in GenBank: Tetraspanin-8 (TET8) *Cucumis sativus* Accession XP_004151158.1, Lectin (phloem protein 2) *Cucumis sativus* Accession NP_001292700.1, and ASSVd RNA sequence was of an isolate from Solan with Accession FM178284. The models were generated from the above-mentioned sequences by the Alphafold2 algorithm (Jumper et al., 2021). The quality was assessed using pLDDT scores. The models were visualized using ChimeraX software v 1.7.1 (Meng et al., 2023). The interactions of PP2 with TET8 and trimeric interaction of PP2 and TET8 with viroid RNA was studied using AlphaFold3 (Abramson et al., 2024; https://alphafoldserver.com/fold/20a59528fa980c38) that allows structural modeling and interactions of protein and RNA. The quality was assessed using pTM scores. The models were visualized using ChimeraX software v. 1.7.1 (Meng et al., 2023).

**Densitometric analysis**-Densitometry was carried out using freely available ImageJ software (https://ImageJ.net/ij/)

### EV acquisition and viroid transmission assay

The whiteflies used for the experiment were glasshouse whiteflies (*Trialeurodos vaprorarium*) (Walia et al., 2015) that were maintained on healthy cucumber and tomato plants in an insect-proof cage. For ASSVd acquisition by whiteflies, ten whiteflies were fed each with AWF (depleted of EVs) and EV (depleted of outside RNAs) and water as control. The whiteflies were trapped in 50ml falcon tubes with their top covered with stretched parafilm. The AWF, EV, and water were placed on the parafilm along with food color (turmeric) and covered with a coverslip (Walia et al., 2015). The tubes were kept in a growth rack under 14h light and 10h dark period for 48h and the whiteflies were allowed to feed on the solution. After 48h the five whiteflies were killed using the cold shock method and RNA isolation and RT-PCR for ASSVd detection was done as described earlier. The remaining five whiteflies were made to feed on healthy cucumber plants (five plants) in an insect-proof rack. The whitefly-fed plants were pooled P1 (1+2) and P2 (3+4+5) checked for the viroid presence using RT-PCR method with ASSVd-specific primers as described above.

For transmission assay using viroid-infected EVs and AWF, three cucumber plants each with AWF (depleted of EVs) and EV (depleted of outside RNAs) were sap inoculated along with leaf sap as positive control and water as negative control. The plants were pooled for individual treatment and checked for the ASSVd presence at 14dpi by RT-PCR method as described earlier.

Statistical analysis. All data presented here are representative of at least three biological and technical replicates for each. Statistical analysis details are mentioned in the respective figure legends.

## Results

### ASSVd RNA localizes in the apoplast and EVs of cucumber plants

To investigate the presence of viroid RNA in apoplast and EVs, we used cucumber plants infected with dimeric infectious transcripts of ASSVd (Walia et al., 2014). The apoplastic wash fluid (AWF) was isolated from viroid-infected plants, 3-week post ASSVd inoculation, using an infiltration centrifugation method (Fig. S1; O’Leary et al., 2014). SDS-PAGE analysis of the isolated apoplastic proteins showed the absence of the large subunit of RuBisCO (approximately 55 kDa), indicating that the AWF was free from cytosolic contamination (Fig. S6a). Additionally, trypan blue staining of the infiltrated leaves revealed the absence of any cell damage and potential cytosolic leakage (Fig.S7). Using size exclusion chromatography (SEC), the isolated AWF was fractionated into 23 fractions (Fig. S2). Of these, fractions 1-7 contained the void volume, fractions 8-11 contained EVs, and fractions 12-23 contained larger protein complexes. Fractions 8-11, corresponding to EVs, were pooled together and examined using Transmission Electron Microscopy (TEM) which confirmed the presence of vesicle-like structures at 50nm, 200nm, and 500nm size corresponding to the size of exosomes and micro-vesicular bodies in plants (Rutter et al., 2017) (Fig. 1a). The pooled EV fractions were also subjected to Dynamic Light Scattering (DLS) for particle size measurement, which revealed their size ranging 50-1000 nm (Fig. 1b). Three peaks corresponding to the size 15nm, 100nm, and 500nm were obtained. The percentage distribution for the 500nm size EVs was maximum compared to 100nm size EVs.

**Figure 1:**
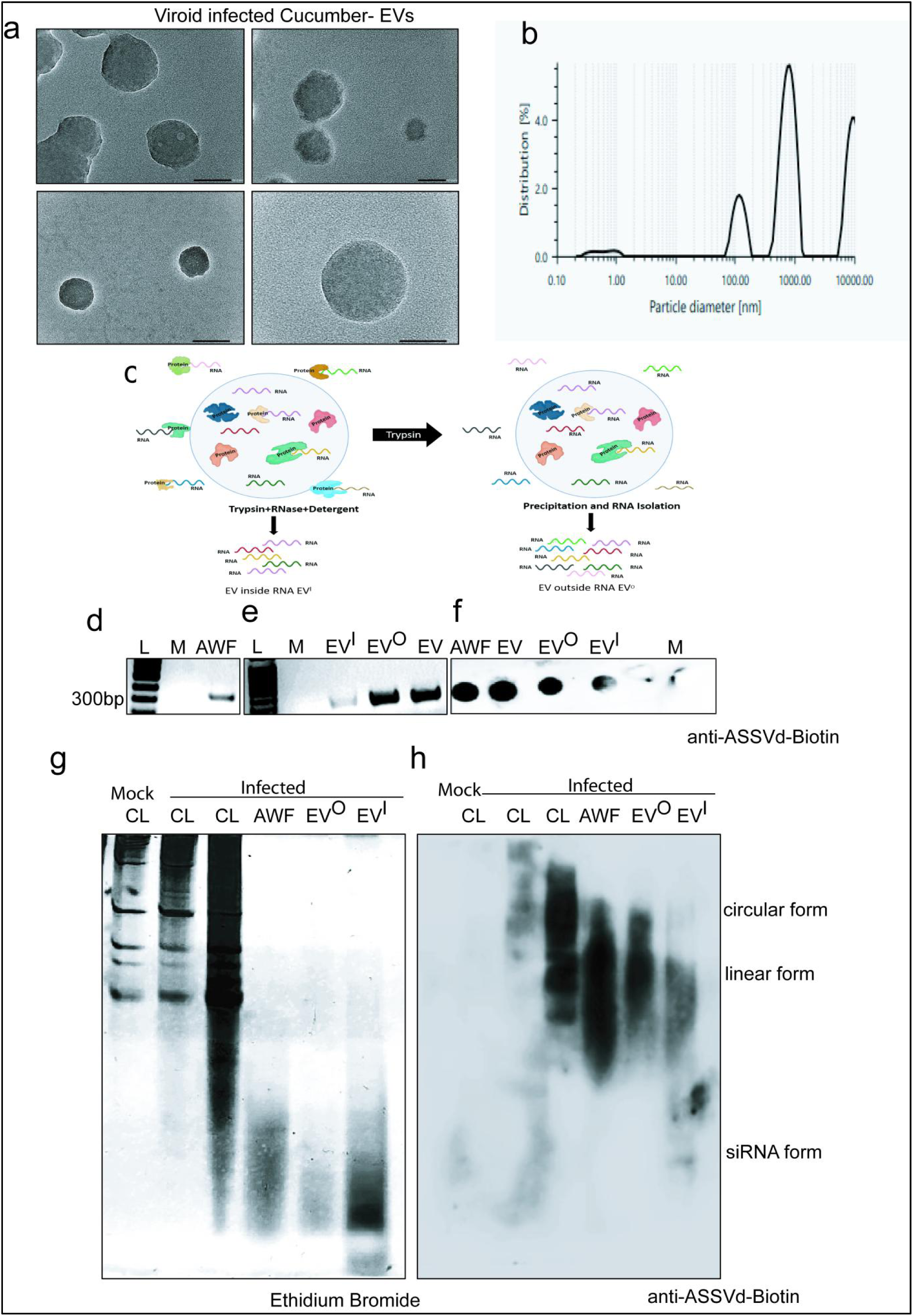
Isolation and characterization of EVs from cucumber apoplast and detection of ASSVd from apoplast and EVs (a) TEM images illustrating the morphology of EVs isolated from cucumber plants infected with ASSVd using the SEC method. EVs are observed to range in size from 50 nm to 500 nm. (b) DLS analysis of the isolated EVs showing particle sizes between 10 nm and 1000 nm, consistent with he sizes observed in TEM. Each experiment was repeated thrice for validation. (c) Graphical method illustrating the protocol followed for isolation of RNA from outside surface of EVs and rom EVs inside. (d, e and f) RT-PCR and dot blot detection of ASSVd in the AWF, total EV fraction obtained via SEC (EV), EV^O^ fraction after trypsin treatment and RNA isolation without EV disruption, and EV^I^ fraction after trypsin, RNase and detergent treatment followed by RNA solation from inside the EVs. (g and h) Northern blot analysis of the RNA samples described above: left panel – EtBr-stained gel showing RNA from infected leaves and their corresponding AWF and EV fractions, with mock leaf RNA as a control; right panel – Northern blot of the same gel using a 3’-labeled biotin-negative sense ASSVd probe, revealing both circular and linear forms of ASSVd in infected leaves, AWF, and inside and outside EVs. Small RNAs of ASSVd are distinctly visible within the EV fraction. (Each experiment was repeated at-least three times for validation)

To examine the presence of ASSVd RNA in the apoplast and EVs, total RNA was isolated from both AWF and pooled EV fractions. Given that RNA can be present either inside EVs or bound to their surface as protein-associated RNA complexes (Karimi et al., 2022), the EVs were further split into two fractions to pinpoint the localization of ASSVd RNA within these vesicles (Fig. 1c). First, the EV fraction was treated with trypsin to release the protein associated RNA and the RNA was isolated from the outer surface of the EVs without breaking the vesicles which represented the RNA associated with the EV surface or EV outside RNA (EV^O^). The remaining EV fraction was treated with trypsin to remove proteins, followed by RNase treatment to degrade protein-bound RNA from the EV surface. The EVs were then ruptured by treatment with detergent and the inside RNA was isolated which represented the RNA inside EVs (EV^I^). The RT-PCR results indicated that ASSVd RNA is present in the apoplast, adhered to the surface of EVs (as EV surface RNA), and is also found inside the EVs (Fig. 1d and 1e). However, the intensity of the ASSVd band was weaker in the inside EV fraction compared to AWF and outside EV fractions as also seen in the dot blot of the same RNA samples (Fig. 1f), suggesting that the mature form of ASSVd RNA is present at lower concentrations inside the vesicles. To further validate these findings, a denaturing Northern blot analysis of total RNA from AWF and inside and outside EV fractions was carried out. The analysis revealed the presence of different forms of ASSVd RNA (including circular, linear, and small RNAs) in both the apoplast and EV fractions. Ethidium bromide staining of the denaturing PAGE gel with total RNA from AWF, EV^O^, and EV^I^ fractions showed an enrichment of small RNAs inside EVs compared to the other fractions, consistent with earlier findings in Arabidopsis (Fig.1g) (Karimi et al., 2022). Furthermore, the Northern blot results confirmed the presence of both circular and linear forms of ASSVd RNA in the apoplast and in both the outer and inner EV fractions, implicating that ASSVd RNA is indeed associated with EVs in these plants (Fig.1h and Table 1). Small RNA species corresponding to ASSVd were detected only within the EVs, suggesting that ASSVd-derived small RNAs (sRNAs) are packaged into these vesicles. The lack of probe hybridization with the RNA from the mock leaf confirmed that the bands observed in the Northern blot were specific to ASSVd and not from any other plant-derived RNAs.

**Table 1:** Representation of northern blot of denaturing PAGE of RNA hybridized with antisense ASSVd probe 3’ labeled with biotin which shows from mock cell lysate, viroid infected cell lysate, viroid infected AWF, outside EVs (EV^O^) and inside EVs (EV^I^)

| Detection Technique | Detection by RT-PCR | Detection by Northern Blot |  |  |
| --- | --- | --- | --- | --- |
| Location/Form | Circular/Oligomeric form | Circular form | Linear form | SiRNA form |
| Apoplast | + | + | + | - |
| Outside EV surface | + | + | + | - |
| Inside EV | + | mild | Not detectable | + |

### ASSVd RNA exhibits characteristics of EV loading including m6A modification and EXOmotifs

An immunoprecipitation (IP) experiment using antibodies specific to m6A with equal RNA from viroid-infected AWF and EVs (Fig. 2a) revealed distinct patterns of m6A modification. When the m6A enriched extracts were probed with biotin labeled negative sense ASSVd probe, it was observed that almost 50 percent RNA in the EVs is modified with m6A modification whereas in the apoplast almost 25 percent ASSVd RNA is in the m6A modified state. The remaining percent is either unmodified or modified with some other kind of modification. Thus, overall through these results, we infer that m6A modification of ASSVd could be one of the factors contributing to ASSVd RNA loading inside the EVs or export into the apoplast. Apart from m6A modification the presence of exosome sorting motifs (EXOmotifs) are also known to transport RNA into the vesicles. For this, we carried out a sequence-specific screening of ASSVd which revealed an enrichment of five EXOmotifs in the ASSVd genome viz. UCCC, CUCC, CCCU, UGUG, and GGUG (Fig. 2b). The UCCC and UGUG motifs lie in the terminal left domain, CUCC and CCCU lie in the central conserved region whereas GGUG lies in the pathogenicity domain of ASSVd (Fig. 2c).

**Figure 2:**
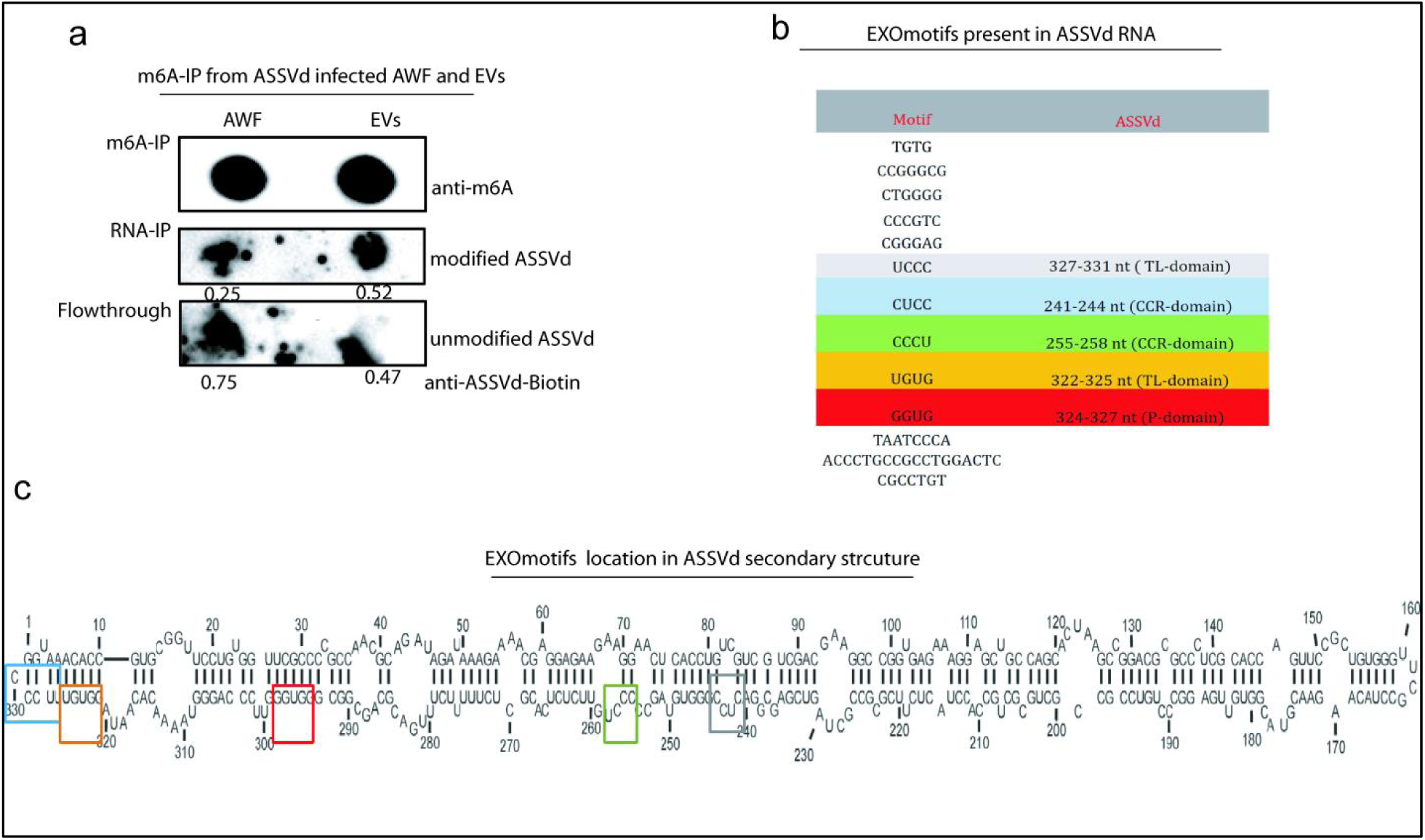
Enrichment of m6A modification and EXOmotifs in the ASSVd genome. (a) Immunoprecipitation of m6A-enriched RNAs from the ASSVd-infected AWF and EVs fractions using an m6A antibody (upper panel), followed by detection of ASSVd in the immunoprecipitated samples using a 3’-labeled biotin-negative sense ASSVd probe (middle panel). The middle panel shows that ASSVd RNA in EVs is modified with m6A, while the lower panel demonstrates that unmodified ASSVd RNA, which did not bind to the m6A antibody, remains in the flow-through fraction. (The assay was repeated twice to confirm the results) (b) Overview of various EXOmotifs identified in animal studies from extracellular RNA and the motifs found within the ASSVd genome. (c) The location of the identified EXOmotifs in the secondary structure of ASSVd is highlighted by colored boxes.

### Investigating protein level changes in the cucumber EVs infected with viroid

For the identification of protein changes in EVs upon viroid stress, EVs from AWF were isolated from ASSVd-infected cucumber plants and mock plants three weeks post viroid infection (Fig. S6b). The morphology of the isolated EVs was analyzed using TEM and DLS which revealed EVs of size varying from 50nm to 1000nm (Fig. S8 and S9). Protein identification of the infected and healthy EVs revealed proteins specific to exosomes such as tetraspanins (Tet16/PLAT/LH2 domain-containing lipoxygenase family protein) in healthy EVs, and lipoxygenase 3 (orthologue of TET8 in cucumber) in infected EVs (Table 2). Additionally, markers for microvesicles, including syntaxins (SYP21/71/31, SNARE-like superfamily protein) and target SNARE coiled-coil domain protein in healthy EVs, and SYP124/21/42/71 and SNARE-like super-family proteins were also found in infected EVs. Proteins associated with EXPO vesicles, such as exocyst complex components (SEC84B, SEC15B, SEC5, EXO70 family protein in healthy EVs, and SEC10, SEC15B in infected EVs) were also identified. These findings align with the TEM data, which confirmed that EVs isolated through SEC had a size of 50-500nm consistent with the size of exosomes, micro-vesicles, and EXPO vesicles in plants. Notably, enrichment of RNA-binding proteins including Argonaute family proteins (AGO2, AGO3, and AGO7), DEAD-box RNA helicases, sorting nexins, and glycine-rich RNA-binding proteins was also observed in both healthy and infected plant EVs. Apart from these RNA-binding proteins, chaperonins such as heat shock proteins (HSP60/70/81/89.1/70, DnaJ) were also identified in the EVs (Table 2). Interestingly, a phloem protein-PP2 was also identified in the cucumber EVs which is known to be involved in viroid transmission to whiteflies (Walia et al., 2015) and contains a dsRNA binding motif. The absence of markers for the trans-Golgi network (syntaxin 61, SYP61), late endosomes (Ara6), Golgi bodies (GmMan149), and plasma membrane (LTI6) confirmed the integrity of the EVs and EV-specificity of the identified proteins (Table 2, File: S1 and S2). Gene ontology (GO) enrichment analysis based on cellular components revealed that the proteins of both healthy and infected EVs were associated with the endomembrane system, cell-cell junctions, the apoplast, the plasma membrane, and extracellular space, among others (Fig.S10) further confirming the purity of the isolated EVs. In the biological process category, the enriched proteins in healthy EVs were primarily linked to the regulation of Ras protein signal transduction and GTPase-mediated signal transduction, while infected EVs proteins were mainly involved in cytoskeleton-dependent intracellular transport and microtubule-based movement (Fig. S11). A deeper analysis of biological processes uncovered a particularly interesting finding: infected EVs were notably enriched in proteins related to non-coding RNA processing/modification and RNA processing/metabolic processes, a category not observed in healthy EVs.

**Table 2:** Displays the list of proteins identified in mass spectrometry data from both viroid-infected and healthy EVs. It includes plant EV-related proteins such as Syntaxins, Tetraspanins, and Expo proteins. Additionally, RNA-binding proteins and Heat shock proteins, previously reported to be associated with plant EVs, are also included. The table further highlights PP2 proteins identified specifically in the EVs from viroid-infected plants.

|  | Healthy |  | Infected |
| --- | --- | --- | --- |
| Accession No. | Protein name | Accession No | Protein name |
|  | **Syntaxins related** |  |  |
| AT5G16830 | Syntaxin of plants 21 (PEP12 VACULAOR TRANSPORT) | AT1G61290 | Syntaxin of plants 124 |
| AT3G09740 | Syntaxin of plants 71 | AT5G16830 | Syntaxin of plants 21 |
| AT5G05760 | Syntaxin of plants 31 | AT4G02195 | Syntaxin of plants 42 |
| AT5G02280 | SNARE-like superfamily protein | AT5G02280 | SNARE-like superfamily protein |
| AT1G15370 | SNARE-like superfamily protein | AT2G35190 | novel plant snare 11 |
| AT1G16230 | Target SNARE coiled-coil domain | AT1G16230 | Target SNARE coiled-coil domain protein |
|  |  | AT3G09740 | syntaxin of plants 71 |
|  | **Tetraspanin related** |  |  |
| AT3G22400 | Lox5/PLAT/LH2 domain | AT1G17420 | Lipoxygenase 3 (PLAT/LH2 domain) |
| AT1G18510 | Tetraspanin 16 (TET8 RELATED) | AT4G02380 | Senescence-associated gene 21 |
| AT4G02380 | Senescence-associated gene 21 | AT2G29350 | Senescence-associated gene 13 |
| AT2G29350 | Senescence-associated gene 13 |  |  |
|  | **EXPO Related** |  |  |
| AT5G49830 | Exocyst complex component 84B | AT5G12370 | Exocyst complex component sec10 |
| AT4G02350 | Exocyst complex component sec15B | AT2G32750 | Exostosin family protein |
| AT2G28110 | Exostosin family protein | AT4G02350 | Exocyst complex component sec15B |
| AT1G21170 | Exocyst complex component SEC5 |  |  |
| AT5G58430 | Exocyst subunit exo70 family protein B1 |  |  |
|  | **RNA binding proteins** |  |  |
| AT1G69440 | Argonaute family protein | AT1G31280 | Argonaute family protein |
| AT1G31290 | ARGONAUTE 3 | AT1G31290 | ARGONAUTE 3 |
|  |  | AT1G69440 | Argonaute family protein |
| AT3G01540 | DEAD box RNA helicase 1 | AT5G26742 | DEAD box RNA helicase (RH3) |
| AT5G07120 | Sorting nexin 2B | AT5G07120 | Sorting nexin 2B |
| AT1G15240 | Sorting nexin, C-terminal | AT5G58440 | Sorting nexin 2A |
| AT3G23830 | Glycine-rich RNA-binding protein 4 | AT3G23830 | Glycine-rich RNA-binding protein 4 |
|  | **Heat Shock proteins** |  |  |
| AT3G07770 | HEAT SHOCK PROTEIN 89.1 | AT3G13860 | Heat shock protein 60-3A |
| AT1G11660 | Heat shock protein 70 (Hsp 70) | AT1G11660 | Heat shock protein 70 (Hsp 70) |
| AT4G02100 | Heat shock protein DnaJ | AT3G07770 | HEAT SHOCK PROTEIN 89.1 |
| AT3G13860 | Heat shock protein 60-3A | AT4G24280 | Chloroplast heat shock protein 70-1 |
| AT3G12580 | Heat shock protein 70 | AT3G23990 | Heat shock protein 60 |
| AT5G56030 | Heat shock protein 81-2 | AT3G08970 | DNAJ heat shock N-terminal |
|  | **Proteasome components** |  |  |
| AT3G61140 | Rpn7 | AT1G04810 | Rpn2/Psmd1 subunit |
| AT1G77440 | 20S Proteasome beta subunit C2 | AT5G42790 | Proteasome alpha subunit F1 |
| AT1G13060 | 20S Proteasome beta subunit E1 |  |  |
| AT3G13330 | Proteasome activating protein 200 |  |  |
|  | **Phloem Proteins** |  |  |
|  |  | AT1G10155 | Phloem protein 2-A10 |
|  |  | AT1G65390 | Phloem protein 2 A5 |

### CsTET8 is a conserved EV marker protein in cucumber while CsPP2 is a newly identified RNA-binding protein localized within EVs

Mass-spectrometry results showed the presence of tetraspanins in cucumber EVs. We further confirmed the presence of TET8 (CsTET8) in cucumber EVs using immunoblotting. To determine whether Arabidopsis TET8 (AtTET8) antibodies cross-react with CsTET8, we conducted sequence analysis and found that the AtTet8 (AT2G23810) ortholog in cucumber, Tet8/Lipoxygenase domain 3 (Csa1g287020), shares 65.67% homology with AtTET8 (Fig. S12). Additionally, the 18 amino acids used to generate peptide-specific antibodies from the EC2 domain (Phytoab, USA) exhibited ∼90% identity between Arabidopsis and cucumber Tet8, suggesting AtTET8 antibodies would likely recognize CsTET8 (Fig. S13).

To validate this, we transiently expressed CsTET8 (pSITE2CA-TET8 and HApBA-TET8) in *N. benthamiana* and performed immunoblotting with AtTET8 antibodies, confirming that AtTET8 effectively detects CsTET8 in cucumber plants (Figs. S14 and S15). Further western blot analysis using AtTET8 antibodies on cucumber samples detected a ∼30 kDa band in cucumber leaves and AWF, consistent with the expected size of AtTET8 in Arabidopsis (Figs. 3a and 3b). These findings confirm that AtTET8 antibodies can effectively detect CsTET8 in cucumber.

**Figure 3:**
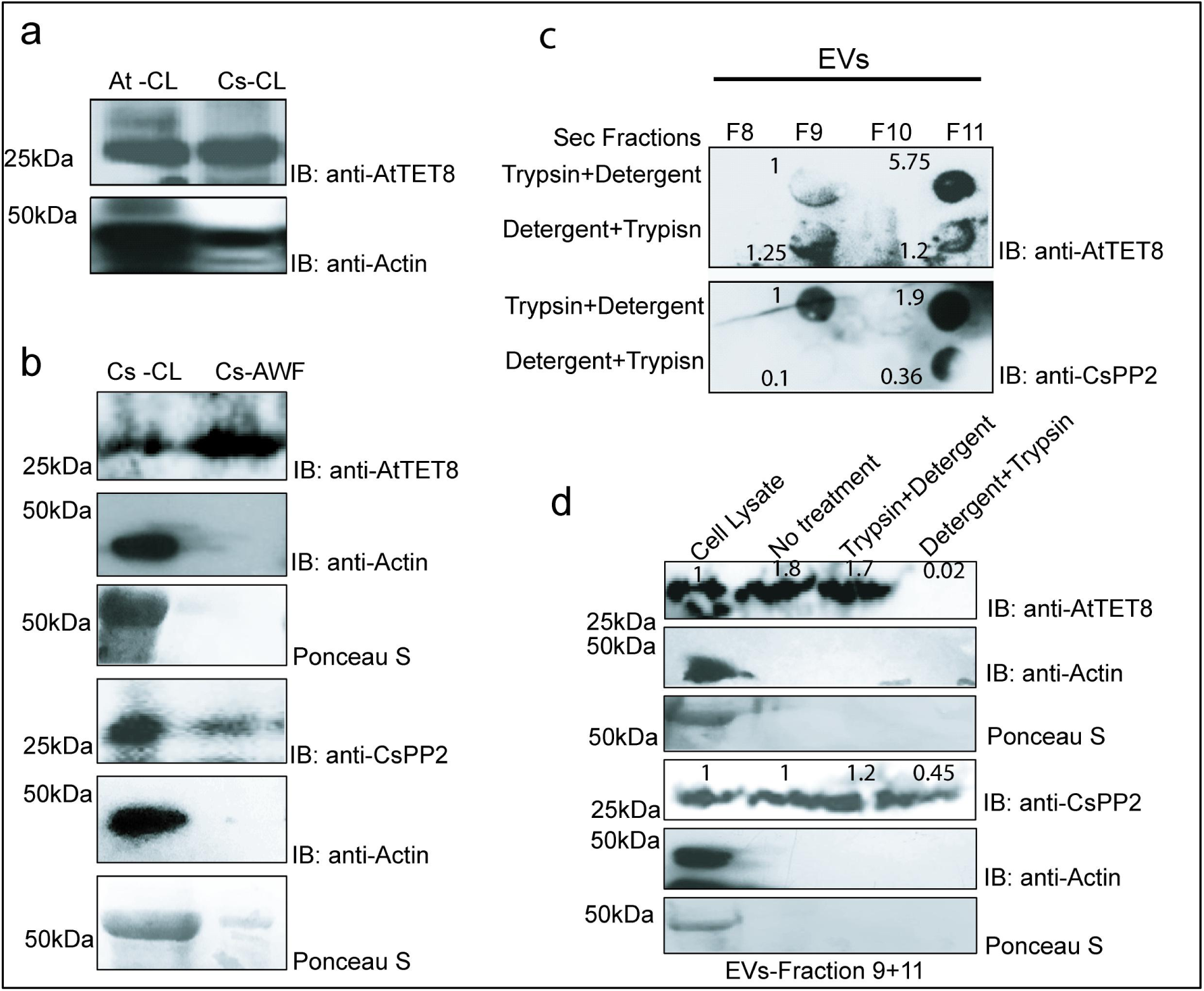
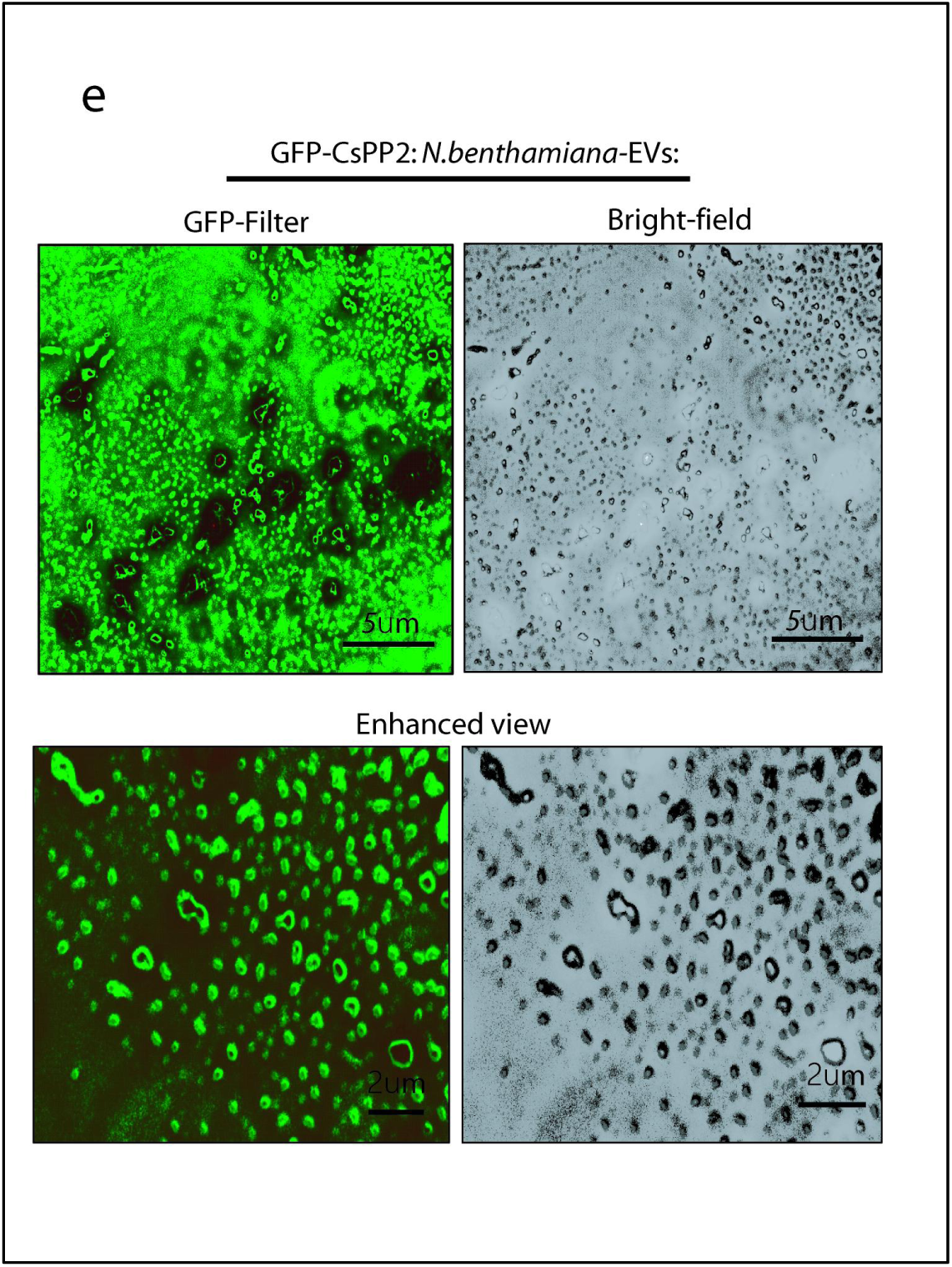
Characterization of CsTET8 and CsPP2 from EVs infected with ASSVd. (a) Western blot detection of TET8 in arabidopsis cell lysate (At-CL) cucumber cell lysates (Cs-CL) using Arabidopsis anti-AtTET8 antibody, revealing a band around 30 kDa, which corresponds to the size of TET8 in both Arabidopsis and cucumber CL (lower panel) western blot of the same extracts with anti-Actin antibody. (b) Western blot of cucumber CL and AWF with anti-AtTET8 and anti-CsPP2 antibody. Western blot of the same extract with anti-actin antibody are used as loading controls. (c)Western blot analysis of EV fractions (8-11) isolated by SEC, using AtTET8 antibodies to detect CsTET8 and CsPP2 in fractions 9 and 11 (upper lane in upper and lower blot). The fractions were treated with trypsin to degrade the surface proteins of the EVs, followed by detergent-induced rupture before loading onto the membrane. The lower lane shows the degradation of CsTET8 and CsPP2 from inside the EVs when ruptured with detergent before trypsin digestion, suggesting both proteins are localized inside the EVs. (d) Western blot analysis using anti-AtTET8 and anti-CsPP2 antibodies of pooled EV fractions (9 and 11) with different treatments—no treatment, trypsin followed by detergent, and detergent followed by trypsin—indicating that CsTET8 and CsPP2 are localized both inside and outside the EVs. Western blot of the same extract with anti-actin antibody are used as loading controls. (Each experiment was repeated three to four times for validation) (e) Fluorescence microscopy of EVs isolated from GFP-CsPP2 expressing *N.benthamiana* plants under GFP Filter and Bright field (Scale bar 5ųm and 2ųm).

To investigate the association of CsTET8 with cucumber EVs, various size-exclusion chromatography (SEC) fractions were analyzed by western blotting. These fractions underwent a protease protection assay: initially treated with trypsin to digest proteins exposed on the surface of EVs, followed by detergent treatment to disrupt the vesicle membrane and release intravesicular proteins. To further confirm the internal localization of specific proteins, EVs were first lysed with detergent and then treated with trypsin, ensuring that only proteins within the vesicles would be degraded.

CsTET8 was detected in EV-containing fractions F9 and F11, with the highest abundance in F11 (Fig. 3c, upper blot, first panel), indicating its presence within EVs. Trypsin treatment alone did not significantly reduce the CsTET8 signal, particularly in F11, suggesting the protein is protected from enzymatic digestion and therefore not exposed on the EV surface (Figs. 3c). However, following vesicle disruption with detergent and subsequent trypsin treatment, CsTET8 bands were degraded, confirming its intravesicular localization (Fig. 3c, upper blot, second panel).

Further validation came from the western blot analysis of pooled EV fractions (F9+F11), which showed a strong CsTET8 signal on SDS-PAGE. This signal was significantly reduced after detergent and trypsin treatment (Figs. 3d), and reflected in the corresponding intensities, reinforcing that CsTET8 is sequestered within the EVs. These findings support that CsTET8 is an integral, protected component of cucumber EVs, consistent with a conserved role across plant species.

Another protein of interest was Phloem Protein 2 (CsPP2) which we proceeded further to explore its association with the EVs. CsPP2 is a phloem lectin previously shown to interact with ASSVd and assist in its cross-kingdom trafficking (Walia et al., 2015). Western blotting of cucumber cell lysate and AWF confirmed the presence of CsPP2 in both intracellular and apoplastic compartments (Fig. 3b). Analysis of SEC fractions revealed that CsPP2 co-eluted with CsTET8 in fractions F9 and F11 (Fig. 3c, lower panel). The protease protection assay demonstrated that CsPP2 also resides within EVs: the intact band persisted after trypsin treatment but was degraded when EVs were pre-lysed with detergent before enzyme exposure (Figs. 3c and 3d). Furthermore, we transiently expressed CsPP2 fused with GFP in *Nicotiana benthamiana* and isolated EVs from the apoplast at 48 hpi. Fluorescence microscopy of the isolated EVs showed a distinct presence of GFP-CsPP2 in the EV fraction, in contrast to mock-treated plants (Fig. 3e and S16). Characteristic small, cup-shaped EV structures expressing GFP-CsPP2 were clearly observed. These findings collectively suggest that CsPP2 localizes both to the apoplast and within cucumber EVs, and when expressed in *N. benthamiana*, it retains the ability to associate with EVs.

Western blot analysis using plant Actin as a control confirmed the absence of Actin protein in both apoplastic and EV samples, verifying the purity of the EV preparations.

### CsPP2 may aid in the sorting of viroid and other plant RNAs into CsTET8-specific exosomes

The co-localization of CsTET8 and CsPP2 in the same EV fractions indicates a potential interaction between these two. To validate this, we carried out immunoprecipitation of CsPP2 using CsPP2 antibody (Walia et al., 2015) from the total cell lysate and AWF (treated with detergent to release the EV proteins) of viroid-infected plants and detected the presence of CsTET8 in the pull-down sample. Our immunoblot analysis revealed the presence of CsTET8 protein in the CsPP2 immunoprecipitated from the cell lysate as well as extracellular fluid (Figs. 4a and 4b). To further validate the interaction, we performed a co-immunoprecipitation assay by transiently co-expressing CsTET8 fused to GFP and CsPP2 fused to an HA tag using the binary vectors pSITE2CA and HApBA, respectively. Immunoprecipitation of GFP-CsTET8 with anti-GFP-conjugated beads resulted in a clear co-enrichment of HA-CsPP2, as detected by immunoblotting with an anti-HA antibody, thereby confirming the interaction between the two proteins (Fig. 4c-upper panel). The GFP pull-down alone did not immunoprecipitate the HA-PP2 (Fig. 4c-lower panel). Additionally, Bimolecular Fluorescence Complementation (BiFC) assays revealed that the interaction occurs both in the cytoplasm and at the plasma membrane (Fig. 4d). Interestingly we observed a punctate-like appearance near the plasma membrane indicating the interactions between the two at plasma membrane (Fig. 4d). These punctate like appearance were also observed when GFP-TET8 and GFP-PP2 were individually transiently expressed in *N. benthamiana* plants (Fig. S17). In addition, the AlphaFold3 tool predicted a pTM score of 0.47 (close to a good structure score of 0.5) for the CsP2-CsTET8 interaction (Fig. 4e). The CsTET8 ortholog in Arabidopsis, TET8_ARATH, contains an extracellular domain (EC2) spanning amino acids 97-235 and the CsPP2 likely interact with CsTET8 at its EC2 domain (Fig. 4e). Overall, together these results validate the interaction between CsTET8 and CsPP2 in the cucumber cell and apoplastic space.

**Figure 4:**
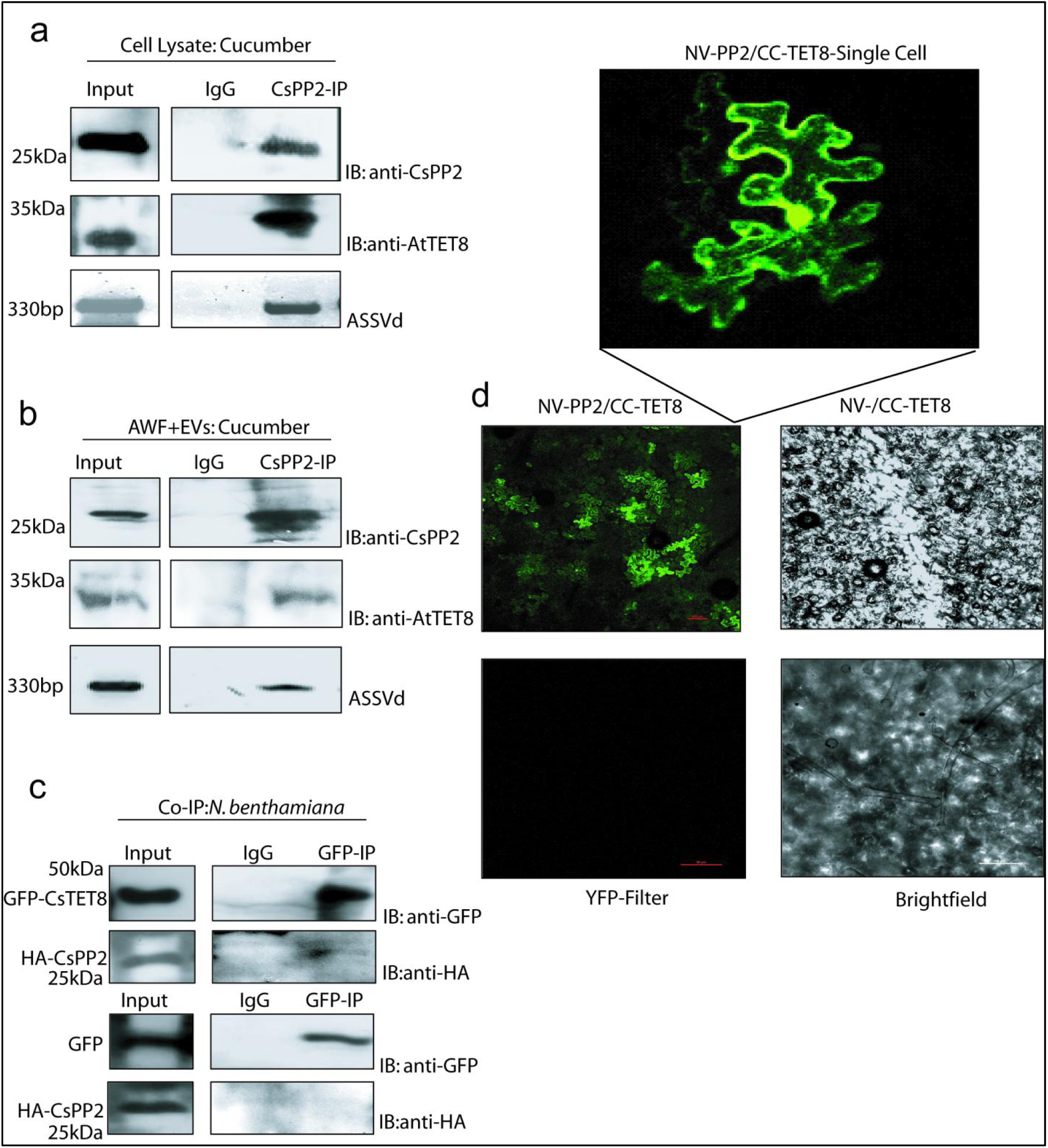
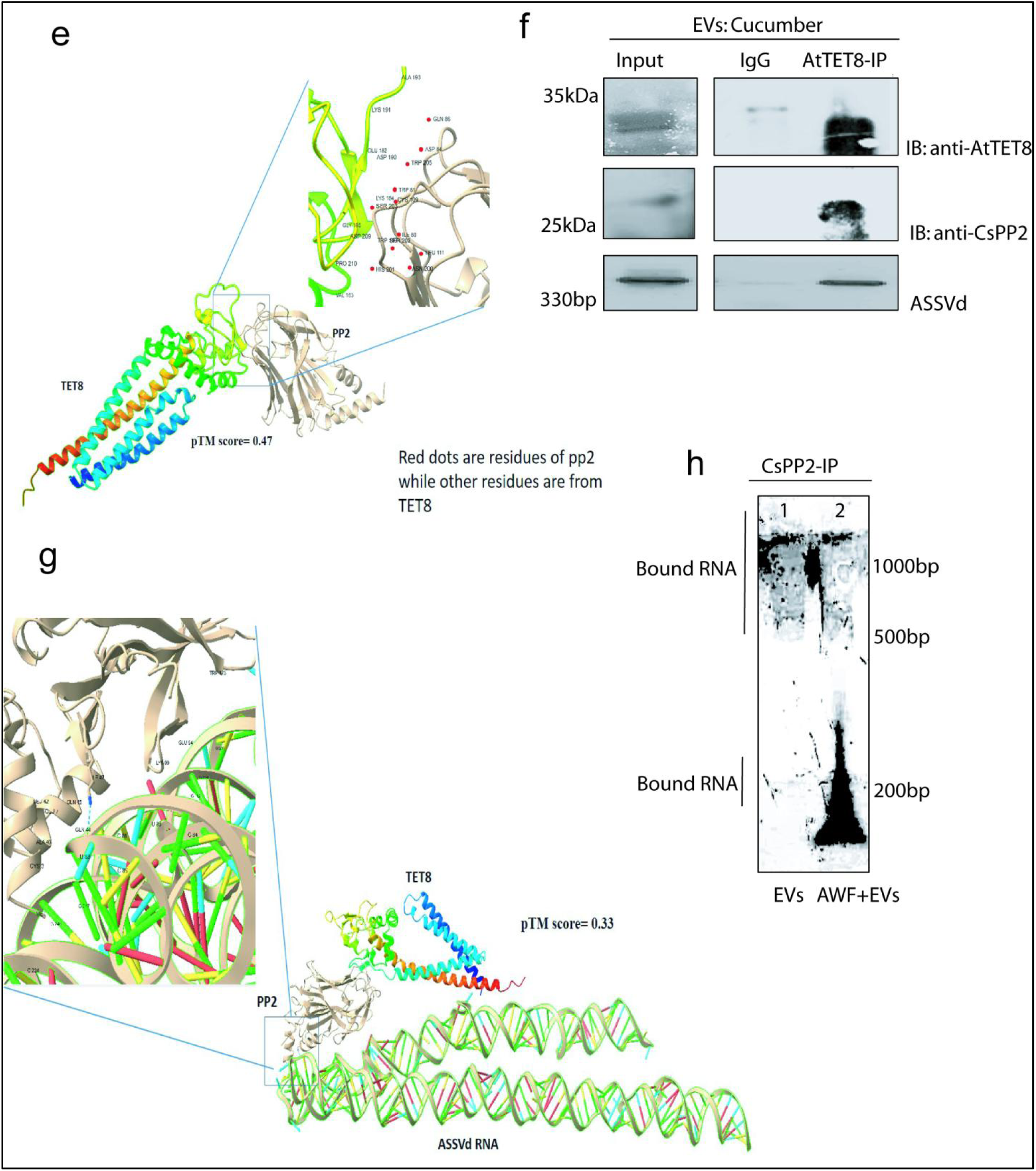
Interaction between CsPP2-CsTET8 and ASSVd **(a)** Western blot analysis showing detection of CsPP2 in its immunoprecipitates from cell lysates (upper panel), along with co-detection of CsTET8 by western blot and ASSVd RNA by RT-PCR from the same sample. **(b)** Western blot showing CsPP2 immunoprecipitated from the apoplast + EVs fraction, with CsTET8 detected via western blot and ASSVd RNA via RT-PCR from the same sample. **(c)** Western blot of a co-immunoprecipitation assay demonstrating the interaction between GFP-TET8 and HA-PP2 when transiently expressed in *N. benthamiana*, using anti-GFP and anti-HA antibodies. **(d)** BiFC images showing the interaction between CC-CsTET8 and NV-CsPP2 observed under a YFP filter. Controls include CsTET8 and NV alone. Bright-field images are shown alongside. A magnified image highlights punctate interaction near the plasma membrane in a single cell. **(e)** Structural prediction of the CsTET8–CsPP2 interaction using AlphaFold 3, with a pTM score of 0.47, suggesting a structure close to the true model. **(f)** Western blot showing immuno-enrichment of CsTET8-containing exosomes from the total EV pool, with co-detection of CsPP2 in the same sample. ASSVd RNA was also detected in the immuno-enriched exosomes using RT-PCR. **(g)** The predicted interaction model of CsTET8, CsPP2, and ASSVd using AlphaFold 3, with a pTM score of 0.33, indicating a model approaching structural accuracy. RT-PCR confirmed the presence of ASSVd in infected samples. **(h)** Total RNA extracted from CsPP2 immunoprecipitates was analyzed on a denaturing gel. The right lane 1 represents RNA from leaf tissue and lane 2 from the apoplast + EVs fraction. Each experiment was repeated at least three times to ensure reproducibility.

**Fig. 5:**
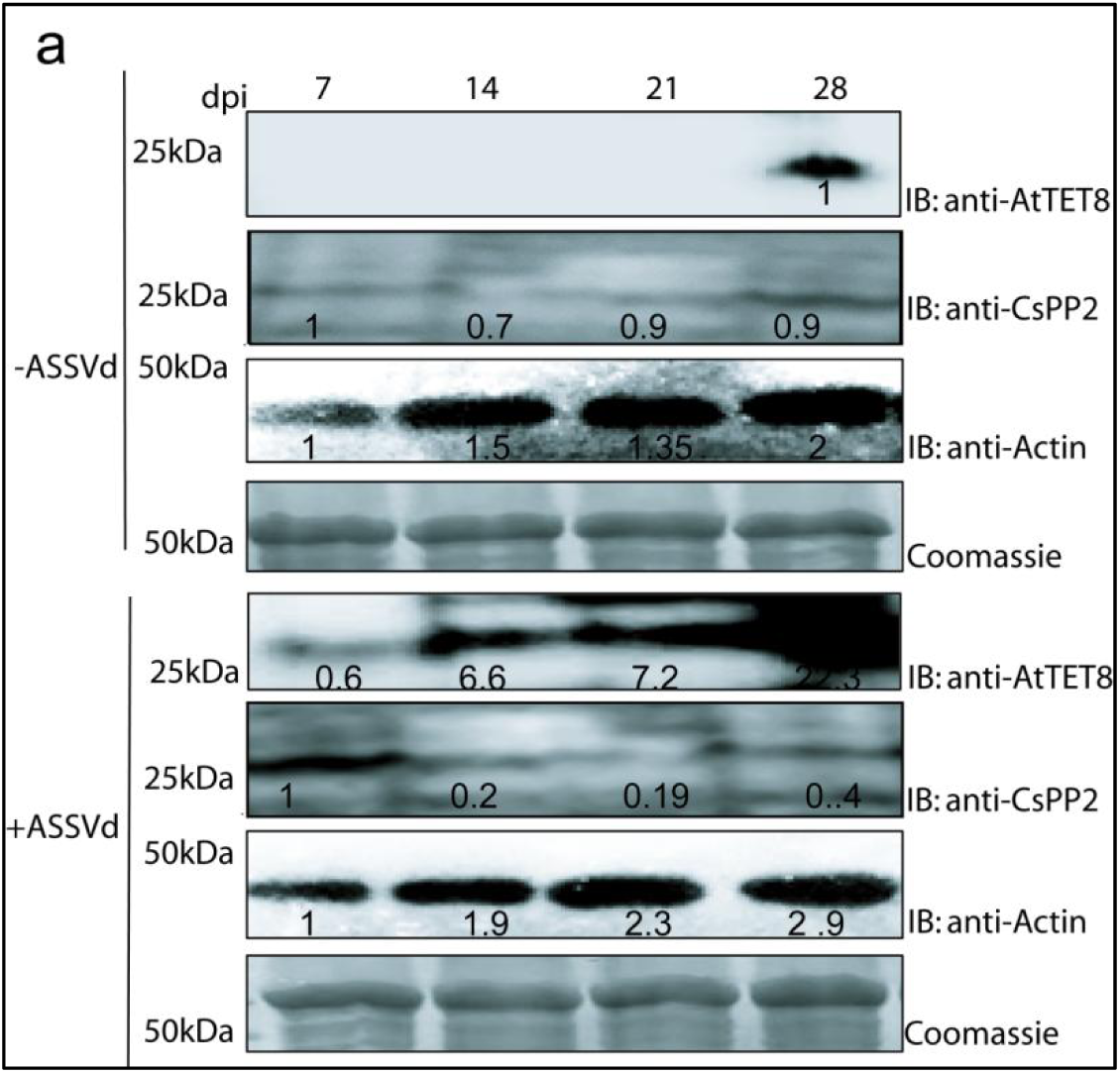
Dynamics of CsTET8 and CsPP2 upon viroid stress (a) Western blot analysis of mock-inoculated and viroid-infected cucumber plants at 7, 14, 21, and 28 dpi, showing the expression levels of CsTET8, CsPP2, and Actin proteins. Blots were probed with anti-CsPP2, anti-AtTET8, and anti-Actin antibodies. Coomassie staining of the same extracts was performed to confirm equal protein loading across samples.

Considering that CsPP2 is an RNA-binding protein, the interaction between CsTET8 and CsPP2 suggests that CsPP2 may play a role in sorting RNAs into CsTET8 exosomes. To explore this interaction, total RNA was isolated from CsPP2-immunoprecipitated samples and the presence of ASSVd RNA was detected by RT-PCR. The results confirmed the co-immunoprecipitation of ASSVd RNA along with CsPP2 from the total cell lysate and apoplast (Fig. 4a and Fig. 4b). Hence, these results suggested that CsPP2, ASSVd, and CsTET8 might exist as a trimeric complex in cucumber plants. To get a deeper insight into this interaction we specifically carried out immuno-enrichment of CsTET8 labeled exosomes from the total EV pool as reported in the arabidopsis (He et al., 2021). Following enrichment, the exosomes were disrupted using detergent and subjected to immunoblotting and RNA extraction. Immunoblot analysis of the enriched CsTET8 exosomes revealed the co-presence of CsPP2, while RT-PCR of the same extracts confirmed the presence of ASSVd within the CsTET8-specific exosomes (Fig. 4f). Interestingly, when the enriched CsTET8 exosomes were analyzed without detergent-mediated disruption, immunoblotting still detected CsPP2, but RT-PCR failed to detect ASSVd. This suggests that ASSVd resides within the exosomes and that there is no direct interaction between CsTET8 and ASSVd (Data not shown). We further validated this model through interactions using Alphafold3, which revealed the tripartite interaction, in which CsPP2 interacts with the viroid RNA on one side and CsTET8 on the other (Fig. 4g). However, CsTET8 did not show any direct interaction with the viroid RNA. The model predicted the central conserved region (CCR) of the viroid RNA, spanning nucleotides 81-96, was in closest proximity to CsPP2 at amino acids 1-8, 41-47, and 91-101, with CsPP2 motifs spanning amino acids 41-47 and 91-101 containing known double-stranded RNA-binding motifs (dsRBM). Additionally, CsTET8 protein motifs 186-193 and 208-211 were in close proximity to CsPP2 regions 78-86, 107-111, and 202-206, which lie within the EC2 of CsTET8 (Fig. 4g). Thus, using a combination of biochemical and bioinformatic tools, we demonstrate that CsTET8 specific exosomes harbor ASSVd and CsPP2 where CsPP2-CsTET8 directly interact with each other at EC2 domain of CsTET8 but there is no direct interaction between ASSVd-CsTET8 further supporting the notion that CsPP2 might facilitate loading of RNAs in CsTET8 specific exosomes. To further demonstrate that CsPP2 could be a potential protein facilitating the sorting of RNAs inside EVs the CsPP2-immunoprecipitated RNAs were separated using denaturing PAGE and stained with Diamond nucleic acid stain, a binding between CsPP2 and RNAs of various sizes was observed in both whole cell lysate and the apoplast/EV fractions (Fig. 4h)) further strengthening the role of CsPP2 as a RNA sorting protein inside vesicles.

### Dynamics of CsPP2 and CsTET8 expression during viroid infection

The expression of CsTET8 is known to be induced upon fungal infection in Arabidopsis (He et al., 2021), while PP2 has been associated with plant defense (Zhang et al., 2011). To explore the role of CsPP2 and CsTET8 upon viroid infection, we monitored the expression dynamics of CsTET8 and CsPP2 proteins in plants infected with the ASSVd. Western blot analysis revealed a gradual increase in CsTET8 expression from 7 to 28 dpi in ASSVd-infected plants, whereas minimal expression was observed in mock-inoculated plants throughout the 21 days (Fig.6a and 6b). However, in viroid-infected plants, CsTET8 expression was elevated as early as 14 dpi, indicating that ASSVd infection triggers the upregulation of CsTET8 at an earlier stage compared to mock-infected plants. In parallel, immunoblot analysis of CsPP2 protein showed a slight increase in expression following viroid infection, peaking at 7 dpi, before returning to normal levels at 14 dpi and remaining constant throughout the infection period (Fig.5a). Hence, both CsTET8 and CsPP2 respond to viroid infection but their expression dynamics differ from each other.

### EV-encapsulated ASSVd RNA is efficiently delivered via mechanical inoculation and is readily acquired and transmitted by whiteflies

To identify the role of EVs in viroid transmission, we conducted an experiment where a group of ten whiteflies (*Trialeurodes vaporariorum*) were fed for 48 hours with three different treatments: intact EVs, AWF (apoplast with EV fractions depleted), and water, in addition to an artificial diet (Walia et al., 2015). After the feeding period, five whiteflies were analyzed for ASSVd presence using RT-PCR, while the remaining five were allowed to feed on healthy cucumber plants. The RT-PCR results revealed that whiteflies were able to acquire ASSVd from the intact EVs (Fig. 6a). Interestingly, a form of infectious ASSVd was also present in the apoplast that had been depleted of EVs, and whiteflies were able to acquire it from there as well (Fig. 6a). However, the PCR signal was significantly stronger for RNA acquired from intact EVs compared to that from the apoplast, indicating that the viroid is more stable and protected from RNases when encapsulated within EVs. This suggests that EVs might facilitate more efficient acquisition of the viroid by whiteflies compared to naked RNA. No ASSVd was detected in whiteflies fed with water.

**Figure 6:**
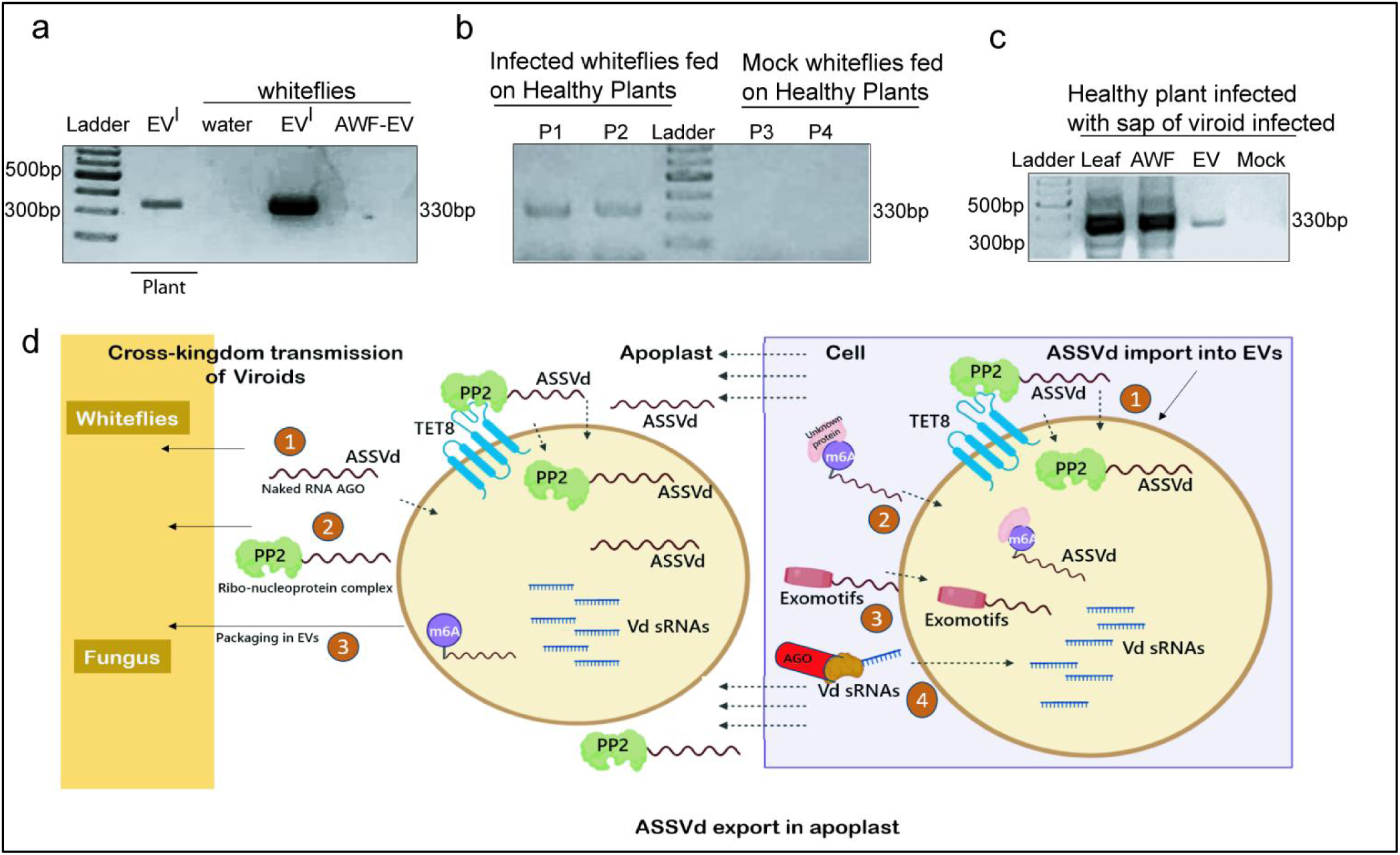
Active transmission of ASSVd to healthy plants and whiteflies via infected EVs. (a) RT-PCR showing detection of the viroid within EVs, whiteflies fed with intact EVs, apoplast lacking EVs, and water as the mock control. (b) RT-PCR showing detection of ASSVd in plants fed with whiteflies that were pre-fed on viroid-infected EVs, with water serving as the control. (c) RT-PCR analysis of cucumber plant sap inoculated with leaf tissue, AWF, and EVs from viroid-infected plants, revealing a 330 bp amplification corresponding to the ASSVd genome using viroid-specific primers. Plants inoculated with water, used as a mock control, did not show the presence of the viroid, serving as a negative control in the assay. (d) A graphical summary of the key findings from the study. In brief, ASSVd RNA (both circular and linear forms) and VdsRNAs can be packaged into EVs. PP2 interacts with ASSVd RNA and the EC2 domain of TET8 within the cell, potentially facilitating the packaging of ASSVd RNAs into EVs. Additionally, ASSVd RNA is modified with m6A inside the cell and contains exomotifs that may aid in its loading into EVs. Once inside the EVs, the viroid RNA is transported to the extracellular space via these vesicles. Viroid RNAs also exist in the apoplast independently of EVs, either as naked RNA species or as RNP complexes. These viroid RNAs can be acquired by whiteflies, and likely by fungi as well, in the form of EV-encapsulated RNA or naked RNAs as RNP complexes, and then transmitted to healthy plants. (The experiment was repeated twice and performed with a set of biological replicates).

Furthermore, plants that were fed with viroid-carrying whiteflies were tested for viroid transmission at 14dpi using RT-PCR, which confirmed that the viroid could be actively acquired and transmitted from EVs through the whiteflies (Fig. 6b). Additionally, we demonstrated that mechanical inoculation of healthy cucumber plants with EVs could also deliver the viroid. For this, we inoculated healthy cucumber plants with intact EVs, AWF, or water. At 14 dpi, RT-PCR on the plants confirmed the presence of ASSVd infection, showing that both the apoplast and EVs of infected plants contained infectious forms of the viroid (Fig. 6c). In contrast, plants inoculated with water showed no viroid-specific amplification. Overall our results suggest that EV-encapsulated viroid RNA can be readily acquired and transmitted by whiteflies to healthy plants.

## Discussion

Viroids, unlike bacteria and fungi, are intracellular pathogens that replicate within the nucleus or chloroplast of the cell. They traffic cell to cell through cytoplasmic connections, plasmodesmata, and through phloem-mediated routes for their long-distance movement. Our previous studies have highlighted the potential role of viroids in cross-kingdom trafficking, such as the acquisition and transmission of ASSVd RNA by whiteflies to neighboring plants (Walia et al., 2015). Later, this phenomenon was also reported in natural interactions between apple plants and their fungal pathogens (Tian et al., 2022). These observations, combined with research on the role of EVs in RNA trafficking during plant-fungal interactions (Cai et al., 2018; Wang et al., 2024), raised important questions about whether the viroid RNAs, typically considered intracellular pathogens, could also be transported in EVs. Our study aimed to investigate the presence of ASSVd RNA in the extracellular space including EVs. Interestingly, our results revealed that viroid RNA also exists in the apoplast, and different forms of ASSVd RNA— circular, linear, and small RNAs—are packaged into the EVs of cucumber plants. This indicated that viroids may employ EVs for *in planta* and cross-kingdom trafficking, as EVs could protect the RNA from host RNases and help viroids cross biological barriers. In animals, certain factors have been identified that assist in the selective loading of RNAs into EVs. For example, certain sequences have been identified in animals that code for the loading of microRNAs into EVs (Martin et al., 2022). These motifs are referred to as EXOmotifs. In search for the identification of such factors that could preferentially facilitate the transport and packaging of viroid RNAs inside EVs, we observed an enrichment of five EXOmotifs in the ASSVd genome. Interestingly, sequence-specific screening of most of the viroids revealed a pattern of exomotif enrichment in their genomes (Fig. S18). For example, the UGUG motif is present in 24 out of 30 viroids, CGGGAG in 2 out of 30, CCGGGG in 16 out of 30, and CCGGUG in 20 out of 30. Notably, the majority of viroids, on average, possess five enriched exomotifs in their genomes. Furthermore, the 25-nt long ACCCUGCCGCCUGGACUCCGCCUGU motif is specifically present in TASVd, which is known to be transmitted by bumblebees through encapsulation in potato leaf roll virus.

Apart from EXOmotifs in animals as well as in plants, the m6A modification of RNA has been linked to the sorting of RNAs into EVs. For example, it has been reported that m6A modification of miRNAs alters their binding affinity to AGO1 protein and increases their association with RNA-binding proteins subsequently facilitating their extracellular export (Sabrina Garbo et al., 2024). In plants, the apoplastic localized circular RNAs and long noncoding RNAs are enriched in m6A which likely facilitates their export into EVs (Karimi et al., 2022). In this study, we found that most EV-localized viroid RNA is modified with m6A however, which of the three forms, whether circular, linear, or small RNA form is modified with m6A still needs to be investigated. Although the biological relevance of this modification is unclear, it is possible that it might have implications on viroid encapsulation within the EVs. Some viroid RNAs are known to undergo reversible modifications, such as m6A methylation (Pavel Vopalensky et al., 2024), thus it is plausible that ASSVd RNA is modified with m6A to enhance its export.

An intriguing finding of our study is the identification of CsPP2, a phloem-specific protein, in the apoplast and EVs of cucumber. A recent study examining the enrichment of the 26S proteasome from melon EVs reported the presence of PP2 in EVs from aphid-infested plants (Sánchez López, 2024). This suggests that the localization of PP2 in the EVs of the *Cucurbitaceae* species could represent a conserved phenomenon. CsPP2 is a phloem lectin with dsRNA binding motifs, capable of increasing the size exclusion limit of plasmodesmata, and has been implicated in the movement of various endogenous and pathogenic RNAs within the phloem (Gomez and Pallas, 2004). In our previous work, we demonstrated that the viroid-PP2 complex is more efficiently acquired and transmitted by whiteflies compared to the naked viroid RNAs (Walia et al., 2015). In the current study, through a feeding assay of EVs to whiteflies, we observed that viroid RNA is acquired and transmitted more efficiently by the whiteflies when packaged into EVs rather than naked RNA or as an RNP complex. This suggests that although viroid RNA can be acquired by whiteflies in various forms—either as a free RNA molecule, an RNP complex or encapsulated within EVs—the transmission through EVs provides a dual advantage: protection from host RNases and a safer route into new host kingdoms. Interestingly, it has been proposed that cross-kingdom trafficking of RNAs can occur via both EVs and RNP complexes. One research group reported that RNA-binding proteins such as AGO1, RNA helicases, and annexins can mediate RNA loading into vesicles and facilitate subsequent cross-kingdom trafficking (He et al., 2021; Cai et al., 2019; Wang et al., 2024). Additionally, another group demonstrated that apoplast-localized proteins like GRP7 and AGO2 can bind to the RNAs in the apoplast, potentially aiding in the cross-kingdom movement of RNAs as RNP complexes (Karimi et al., 2022). Combining these findings, we show using viroid RNA as a model, that cross-kingdom trafficking of RNAs can occur both as RNP complexes and via EVs, although potentially at different intensities. However, the relevance and function of viroid RNAs and CsPP2 within EVs, as well as the potential for viroids to be transmitted through EV encapsulation, require further investigation. The presence of the CsPP2-viroid complex in the apoplast and its association with CsTET8 suggests that CsPP2 may be a novel protein that assists in sorting RNA into EVs in *Cucurbitaceae* plants. Interestingly like TET8, CsPP2 also exhibits the presence of a transmemberane domain in its structure which supports its presence in the extracellular space and EVs (Fig. S19). It is quite possible that viroids can channelize the EV-mediated route for their systemic movement inside plants, as hypothesized for the movement of endogenous RNAs and viral RNAs in plants (Kehr et al., 2018).

The EV proteins from viroid-infected plants implicated changes in nuclear proteins specifically related to non-coding RNA-related biological processes. Viroids are nuclear-replicating pathogens and their effect on nuclear protein turnover and their subsequent export via autophagy or extracellular vesicles warrant further investigation. These findings correlate with cancer disease progression, where genomic DNA and nuclear components are exported by the formation of amphisome-like structures and their fusion to vesicles (Akira et al., 2019). It is quite possible that viroids may influence similar processes during infection, and hence, characterization of DNA and RNA from EVs during viroid infection might shed light on these unexplored processes related to viroid pathogenicity in plants. The induced expression of CsTET8 and CsPP2 proteins suggests that it acts as a positive regulator of viroid stress and could be used as a marker protein for viroid stress. Overall, through our study using ASSVd as the model RNA system, cucumber as its host, and whitefly as its vector, we propose that viroid RNAs are exported into the plant apoplast and packaged into EVs from where they can be acquired by the whiteflies (Fig. 6d). They may use EVs as a means of cross-kingdom trafficking and their interactions with RNA-binding proteins such as CsPP2 could assist their loading into exosomes which improve their transmission efficiency. Moreover, the presence of factors like EXOmotifs or m6A modification may assist viroid RNA in their packaging inside EVs. The precise functional roles of small RNAs in EVs, particularly those derived from viroids and their characterization, remain unclear.

These small RNAs may influence the plant’s immune response or aid in the viroid’s transmission to other organisms, such as insects or fungi. Understanding the mechanisms by which viroids and other RNA molecules are sorted into vesicles and transferred across biological boundaries will offer valuable insights into cross-kingdom RNA trafficking, with implications for viroid biology, RNA-based inter-organism communication, and the ecological impact of RNA transfer.

## Author contributions

**Neha Devi:** Formal analysis, methodology, and validation. **Vasudha Sharma**: Formal analysis and methodology. **Nisha Devi:** Formal analysis and methodology. **Ravi Gupta:** writing-reviewing, and editing (supporting). **Sunny Dhir:** conceptualization, data analysis, investigation, methodology, supervision (equal), writing-review and editing. **Yashika Walia:** conceptualization (lead), data curation and analysis (lead), investigation (lead), methodology, supervision (equal), validation, writing-original draft, writing-review and editing, funding acquisition.

## Acknowledgments

We would like to thank the Management MMDU, for providing necessary research facilities. We would like to acknowledge iSTEM facility at SAIF, IIT Bombay for mass-spectrometry analysis and CIL facility at Panjab University Chandigarh for TEM imaging and Confocal analysis. We would also like to thank I-STEM facility at Jamia Hamdard, New Delhi for confocal microscopy. We acknowledge special thanks to Dr. Vipin Hallan, CSIR-IHBT Palampur for providing CsPP2 antibodies.

## Ethics Statement

not applicable

## Data availability statement

The data that support the findings of this study are available from the corresponding author (YW) upon reasonable request.

## Supporting data

**Fig. S1.**
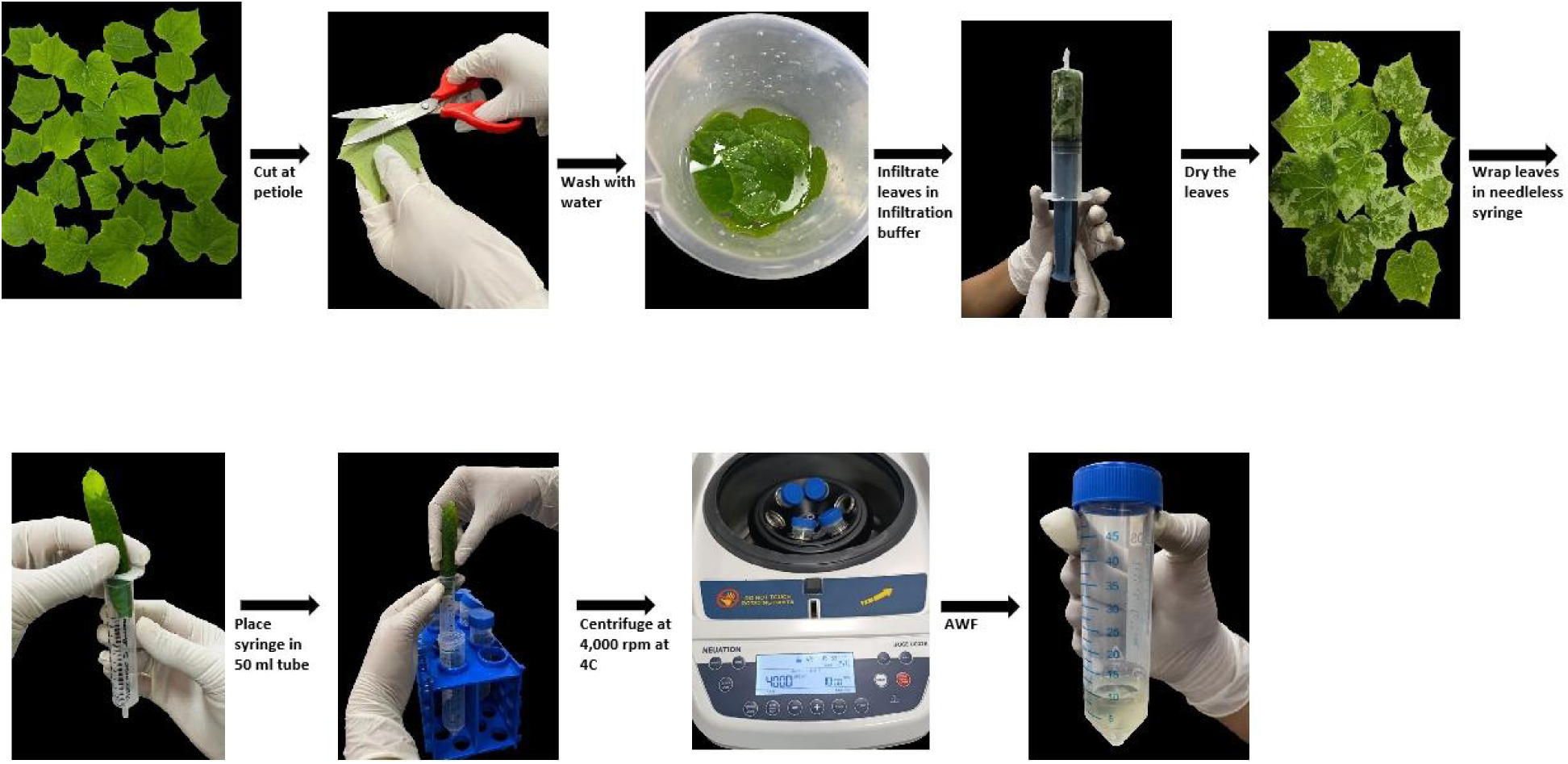
showing the procedure followed for isolation of AWF from cucumber leaves

**Fig. S2.**
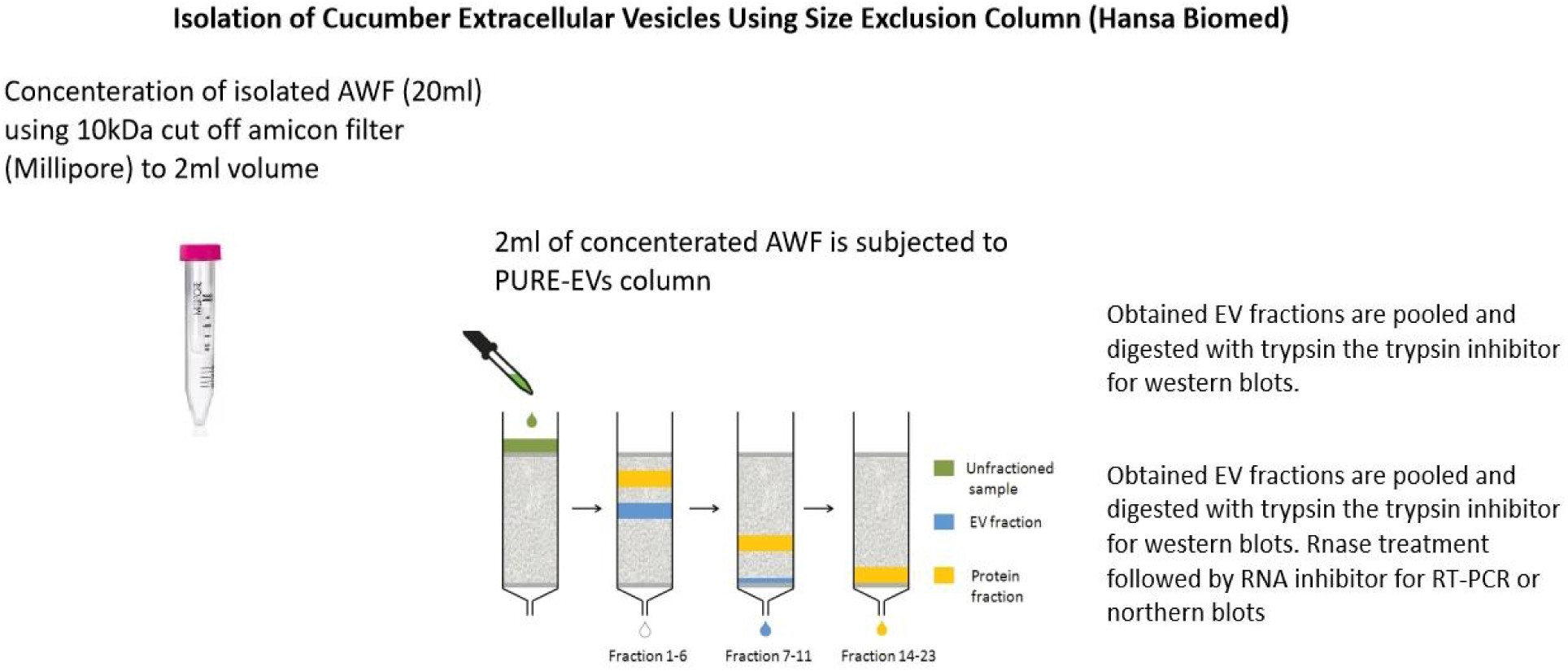
showing the procedure followed for isolation of EVs from isolated AWF of cucumber

**Fig. S3 and S4:**
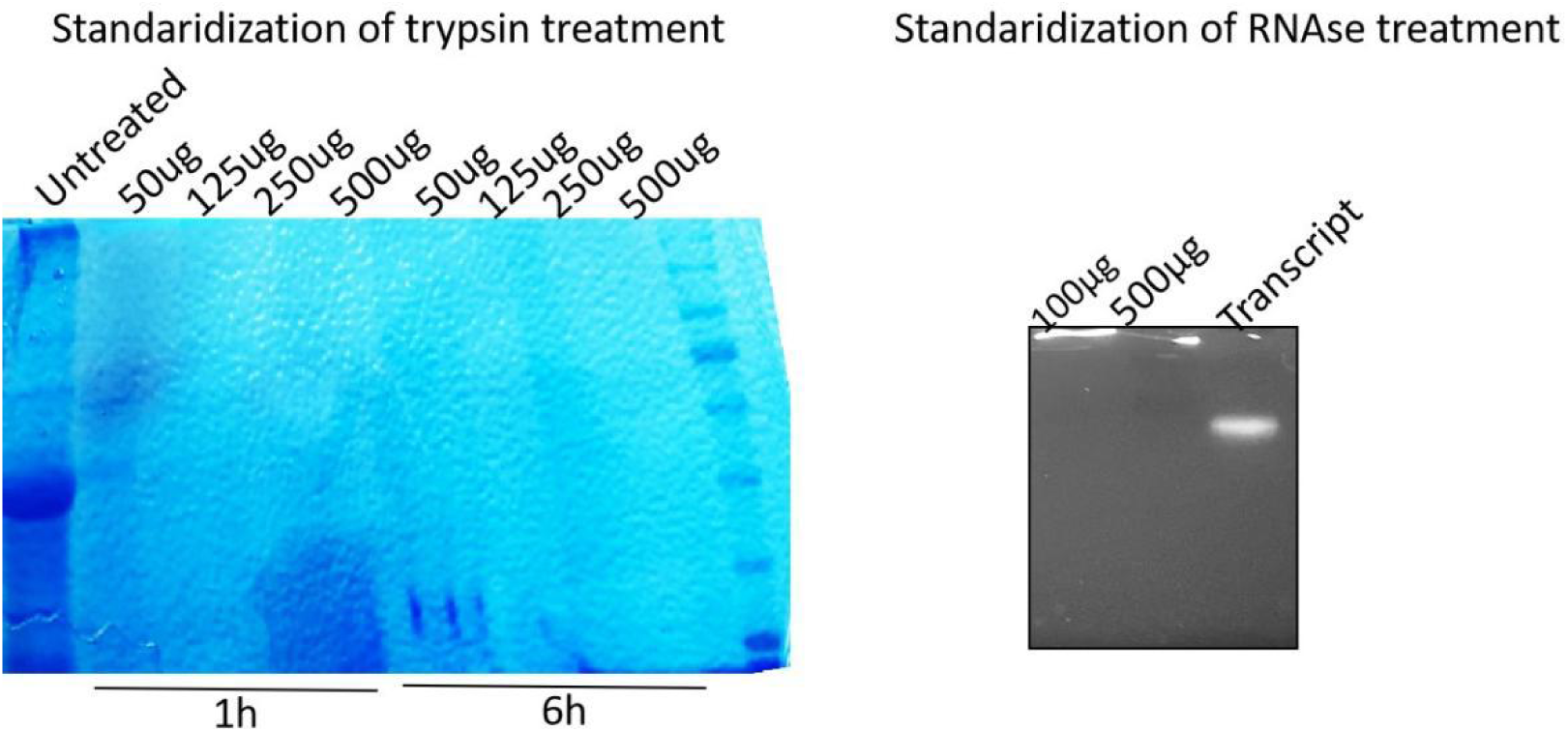
Standardization of trypsin and RNase concentration for treatment of EVs with trypsin and RNase for protease protection assay.

**Fig. S5:**
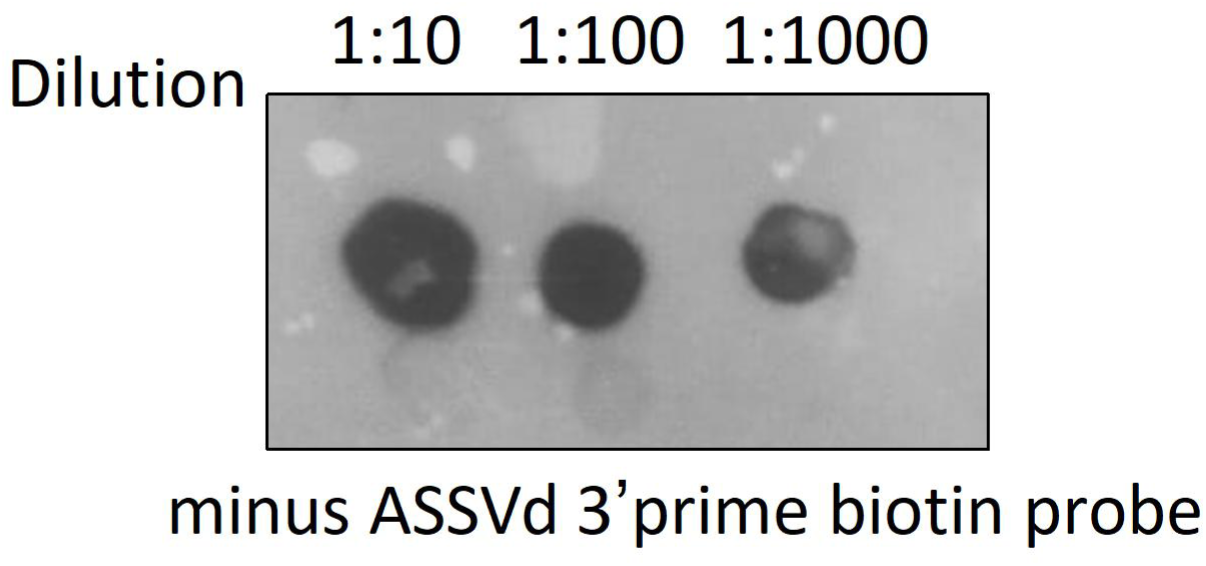
Showing labeling of minus strand of ASSVd transcript with Biotin at its 3’ end and its detection using anti anti-Streptavidin antibody.

**Fig S6.**
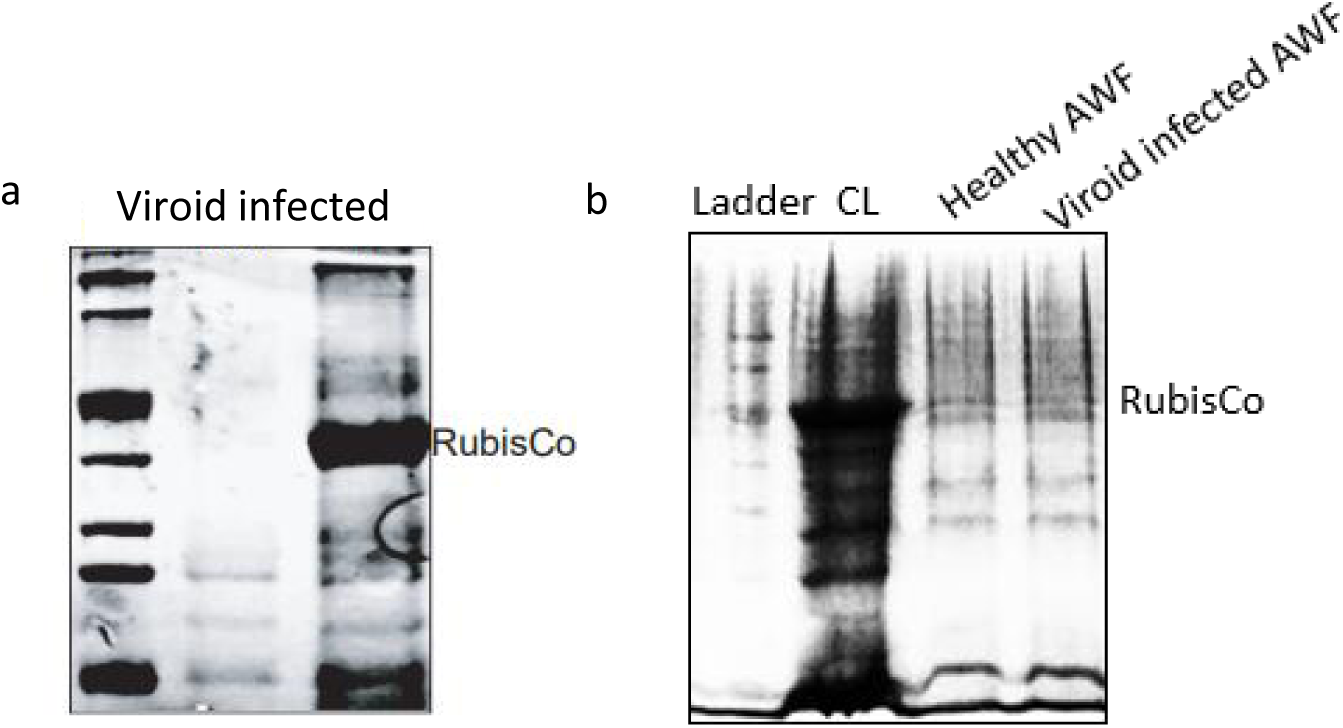
a and b) Coomassie staining of SDS PAGE with total protein extract isolated from AWF and whole cell lysate (CL). Absence of Rubisco protein at 55kDa in AWF fraction compared to CL indicate purity of AWF and absence of cytosolic contamination

**Fig. S7.**
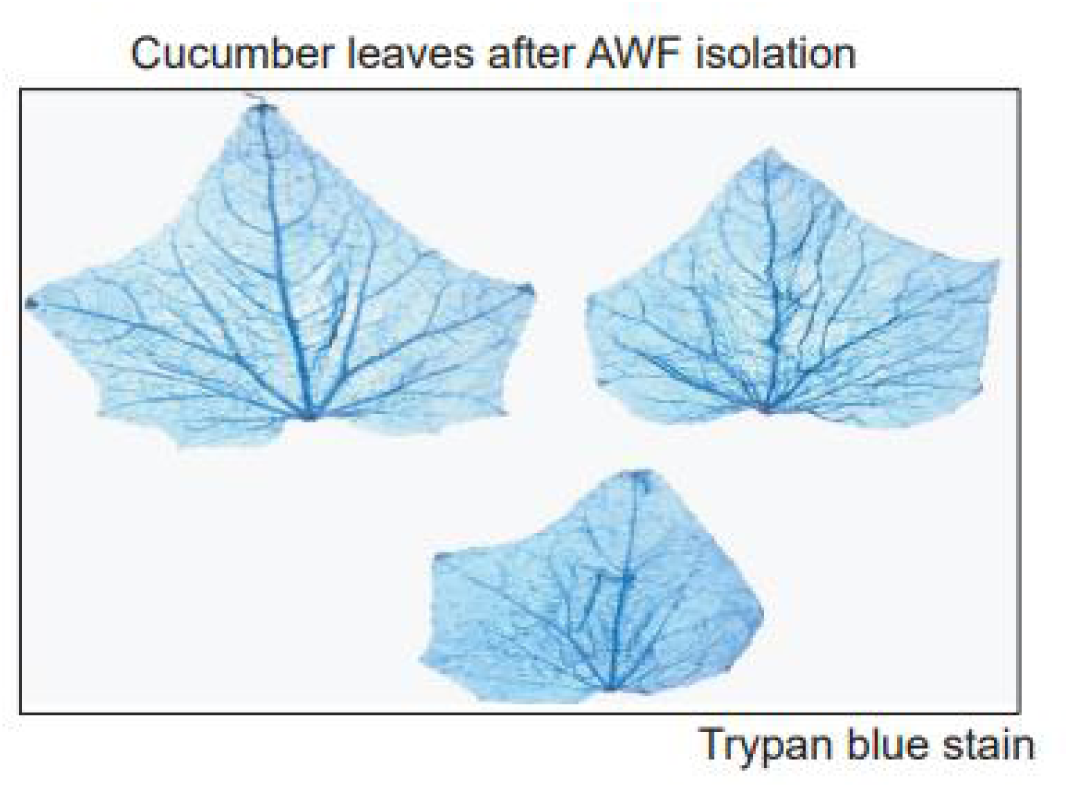
Trypan blue staining of leaves used for AWF isolation. Absence of any cell damage as indicated by clear leaves suggest purity of isolated AWF free from cellular content.

**Fig. S8.**
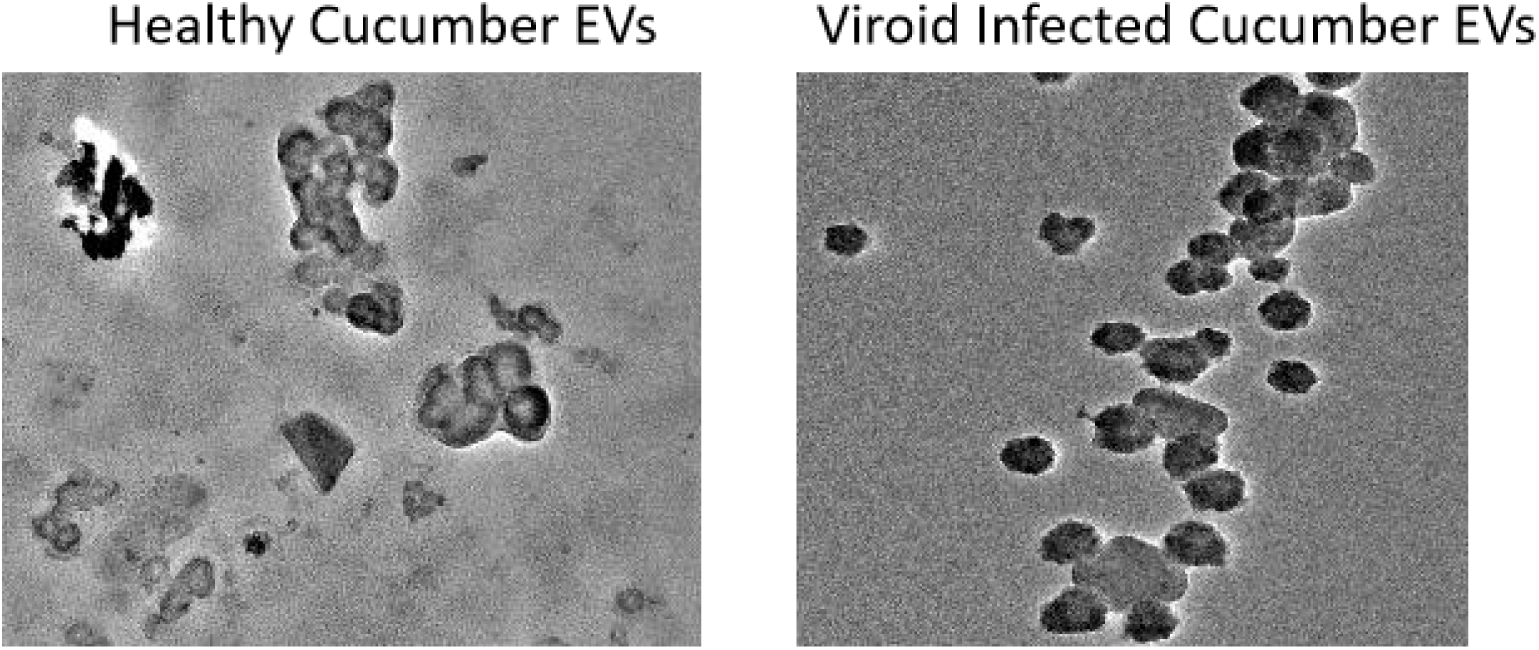
Showing TEM images of EVs isolated from healthy cucumber apoplast and viroid infected cucumber apoplast

**Fig. S9.**
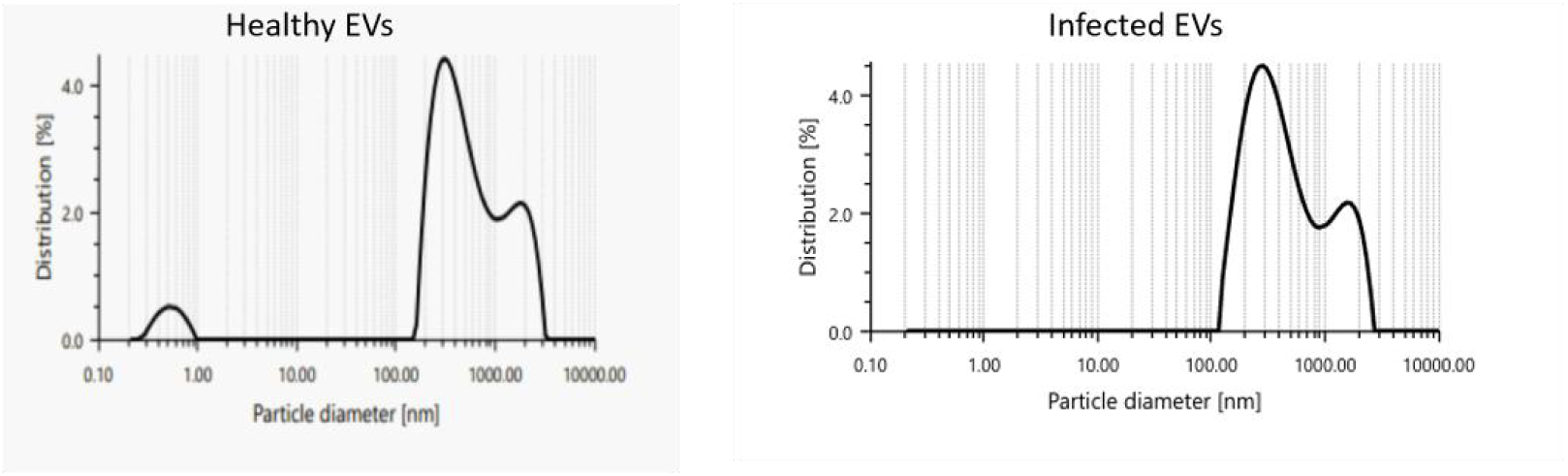
Showing the diameter of particles in nanometre and particle distribution in percentage of EVs isolated from healthy cucumber apoplast and viroid infected cucumber apoplast.

**Fig. S10.**
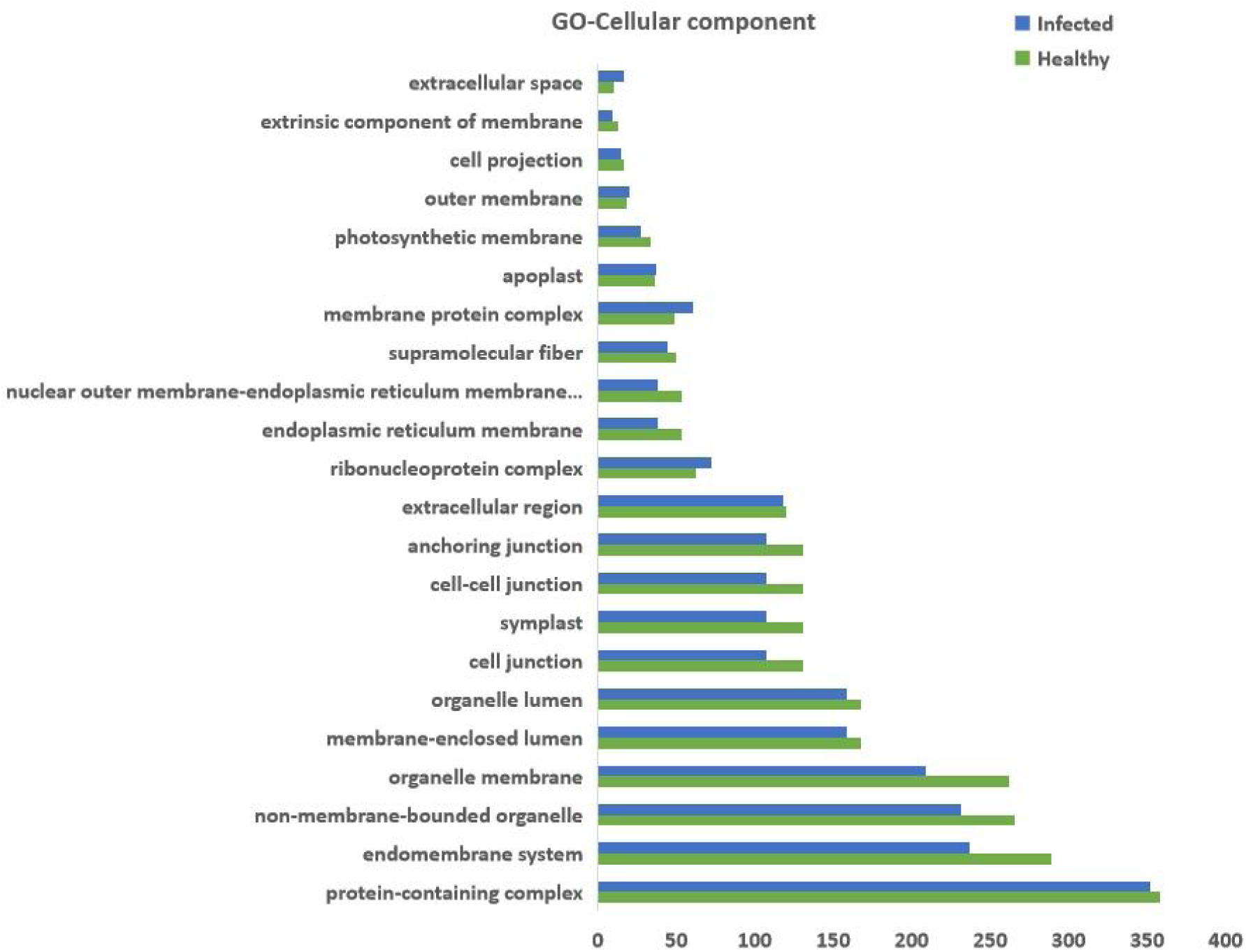
Bar showing classification of viroid infected and healthy EV proteins based on subcellular component

**Fig. S11.**
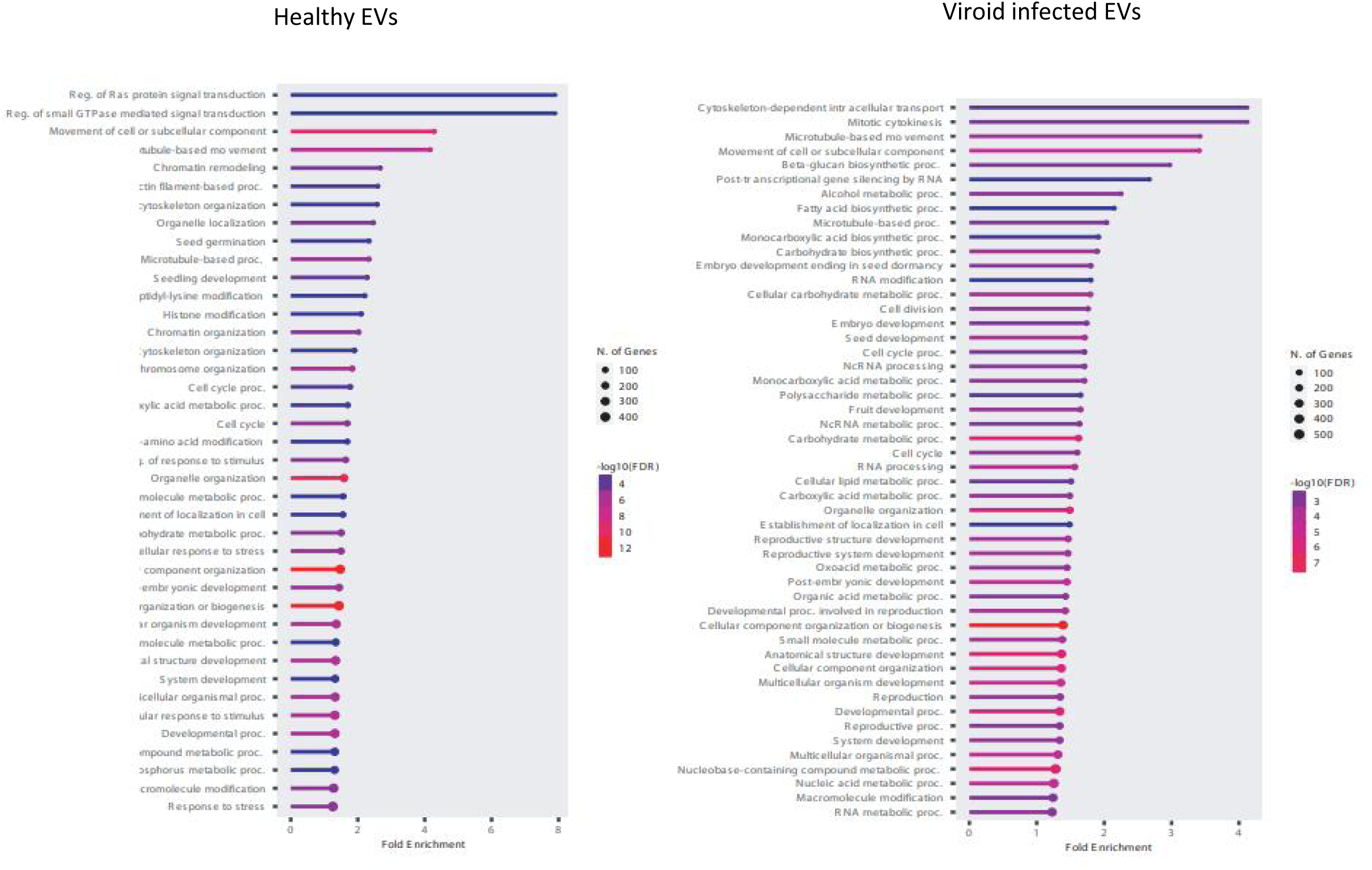
Bar graph showing the classification of viroid infected and healthy EV proteins based on biological processes with notable enrichment of proteins related to non-coding RNA processes in viroid infected EVs compared to healthy EVs. (Each sample is pool of at least 50 biological replicates). We tested whether Arabidopsis TET8 (AtTET8) antibodies would cross-react with cucumber Tet8 (CsTet8) protein and found that the Arabidopsis Tet8 (AT2G23810) ortholog in cucumber is Tet8/Lipoxygenase domain 3 (Csa1g287020) that shared 65.67% homology with AtTet8 (Fig. S7). Additionally, the 18 amino acids used to generate the peptide from the EC2 domain (Phytoab) were over 90% identical between the Arabidopsis and cucumber Tet8, suggesting that AtTet8 antibodies would cross-react with cucumber Tet8 (Fig. S8).

**Fig. S12.**
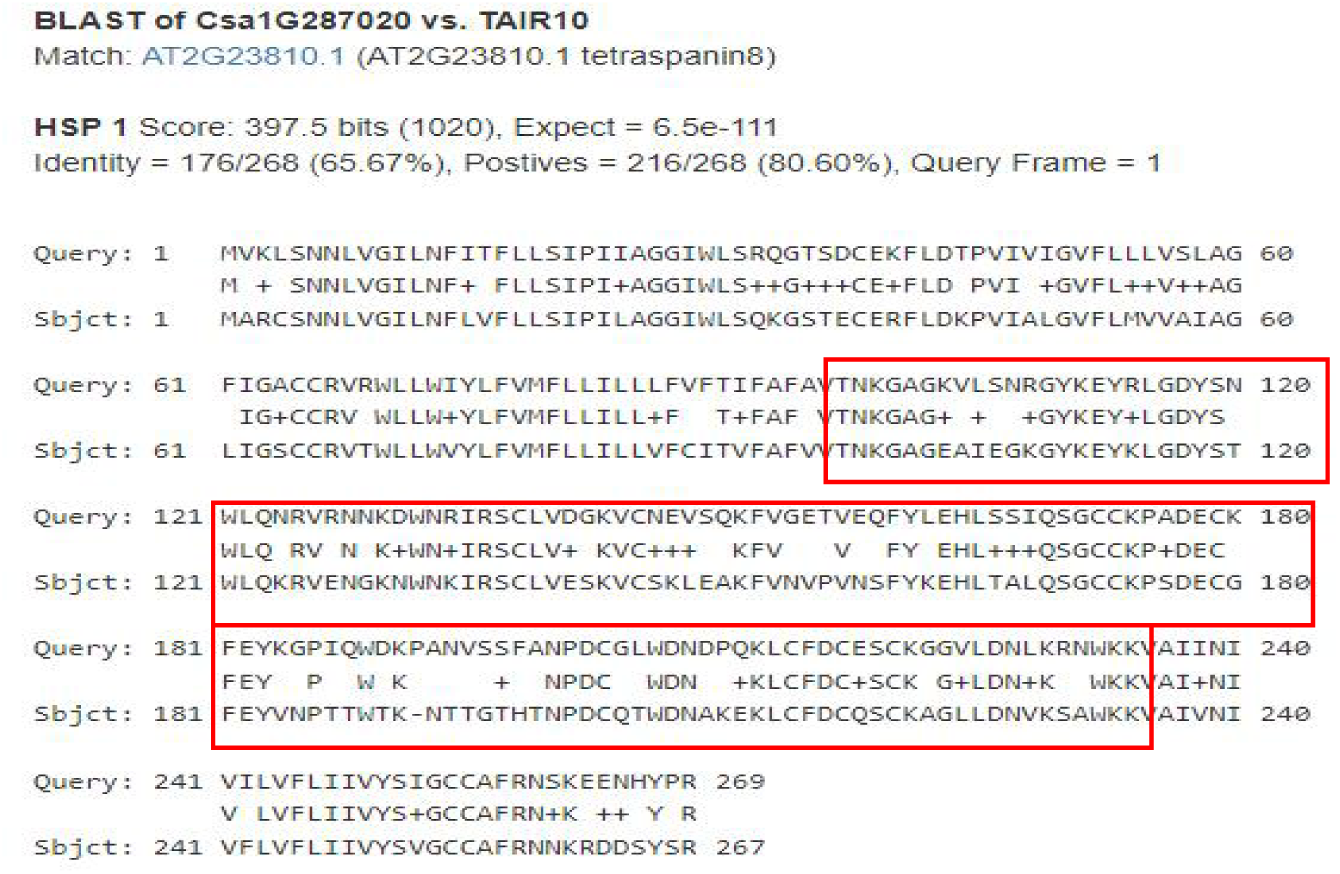
Sequence Homology between CsTET8 and AtTET8 showing 65.67% identity between the two proteins. The EC2 domain froM which the antibody of Phytoab AtTET8 is made is marked in red boxes

**Fig. S13:**
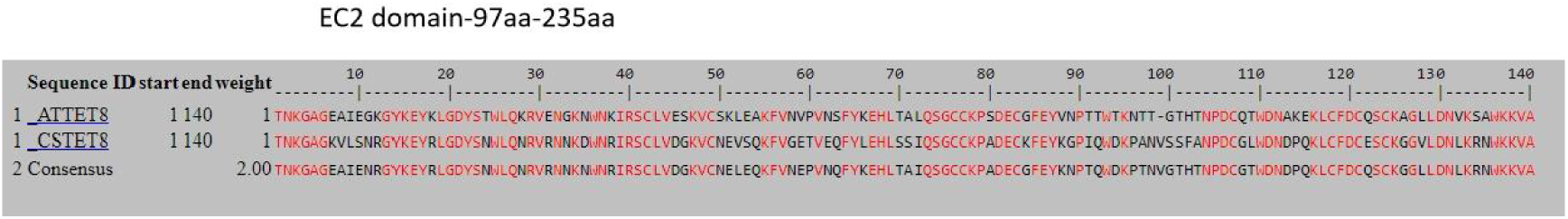
Multiple alignment between EC2 domain of CsTET8 and AtTET8 showing consensus sequences between the two proteins To further confirm this cross-reactivity, we cloned CsTET8 (Csa_1g287020), the cucumber ortholog of AtTET8, into the pSITE2CA and HApBA plant binary vector with a GFP tag for visualization. These recombinant plasmid and were then transiently expressed in Nicotiana plants, and western blot analysis using anti-GFP, anti-TET8 and anti-AtTET8 antibodies confirmed that the AtTET8 antibodies effectively recognized CsTET8 (Fig. S14 and S15) at size of approximately 55kDa and 25kDa, respectively. Based on these findings, we proceeded to use the available AtTET8 antibodies in further experiments.

### Generation of constructs

The CsTET8 gene (Csa1g287020) was amplified from cucumber plants using the RT-PCR method. In brief, RNA was extracted from healthy cucumber plants with RNAiso Plus. Reverse transcription was performed with the Verso cDNA synthesis kit (Thermo). PCR amplification was carried out using Phusion DNA polymerase (Thermo) with specific primers (CsTET8F/R) (Table S1). The PCR products were resolved on a 1.5% agarose gel, and the desired bands were excised and purified using the GeneJET Thermo kit. The purified fragments were then cloned into the pJET1.2 vector (Thermo) and transformed into TOP10 competent cells. The recombinant plasmid was sequenced, confirming 100% identity to CsTET8.

The CsTET8 gene was further amplified using adaptor primers (CsTET8att F and CsTET8att R) and universal primers (Adaptor F/R) (Table S1). The amplified fragment was ligated into the PDONR207 vector using BP clonase (Invitrogen) and transformed into TOP10 competent cells. The transformed cells were plated on LA plates containing 100 mg/ml kanamycin. Recombinant plasmids were verified through colony PCR using the vector-specific primer (Nru R) and the gene-specific primer (CsTET8 F).

Subsequently, the pDONR-CsTET8 clone was sub cloned into the pSITE2CA vector and the HA-pBA vector (Chakrabarty et al., 2007) for expression under the CaMV35S promoter, with N-terminal GFP and HA tags, respectively, using the LR clonase enzyme (Thermo Scientific, USA). The ligated products were transformed into LA plates containing 50 mg/ml spectinomycin. Recombinant clones were screened by colony PCR with the CaMV35S forward primer and CsTET8R primer. The GFP-TET8 and HA-TET8 constructs were then transformed into the Agrobacteriun strain GV3101 using chemical transformation and used for transient expression analysis in *Nicotiana benthamiana* plants, as described by Walia et al. (2020).

**Fig. S14:**
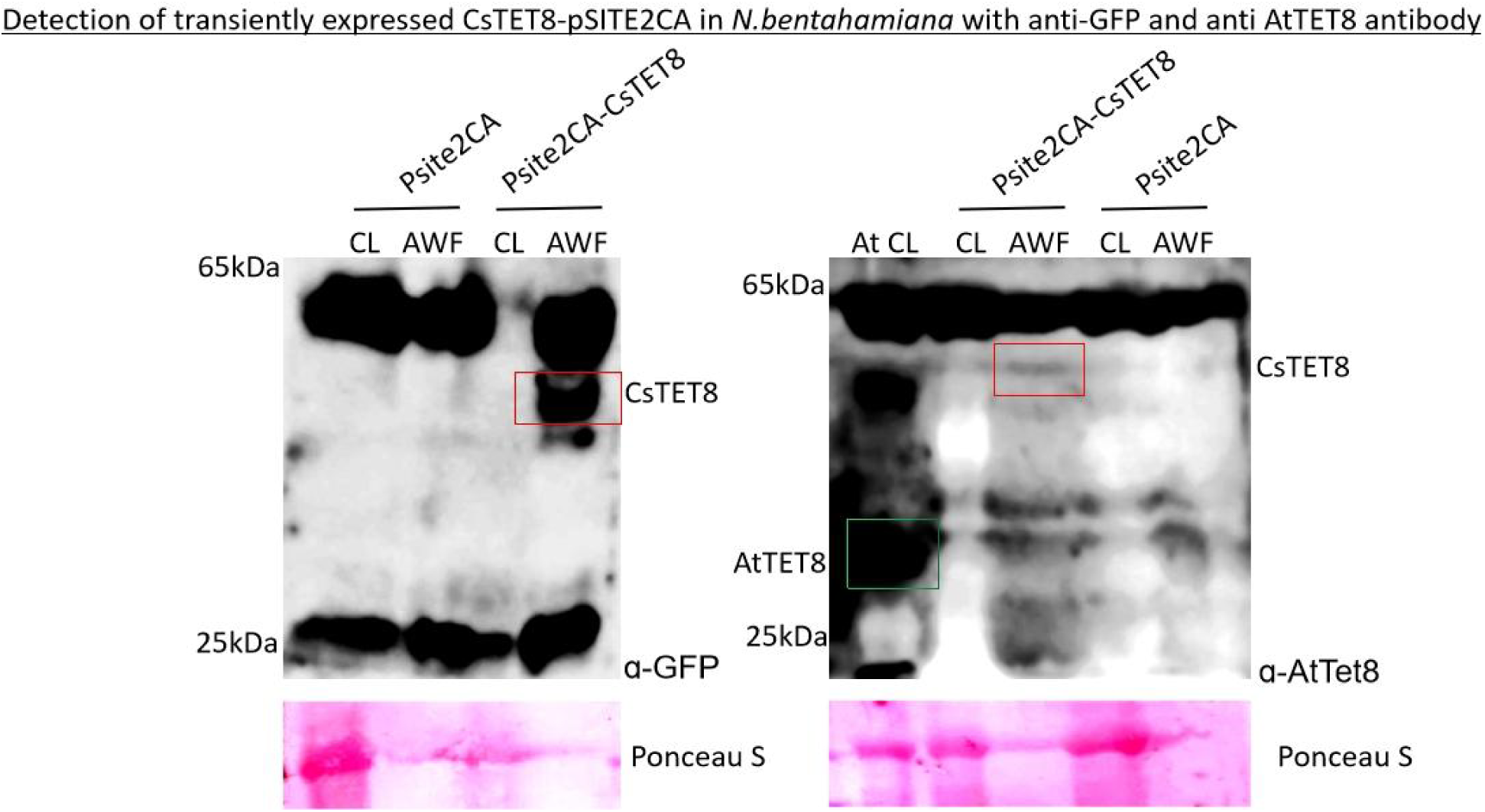
a and b: western blot of *N. benthamiana* cell lysate (CL) and AWF showing transient expression of CsTET8-pSITE2CA in and empty vector (pSITE2CA) with 25kDa band corresponding to the size of only GFP and app. 55kDa size of CsTET8+GFP detected with (a) anti-GFP antibody. (b) AtTE8 antibody. The band corresponding to the size of CsTET8+GFP is detected in AWF and marked in the box. The arabidopsis TET8 band detected with AtTET8 antibody at 30kDa as positive control is marked in the green box (Fig.S9b)

**Fig. S15:**
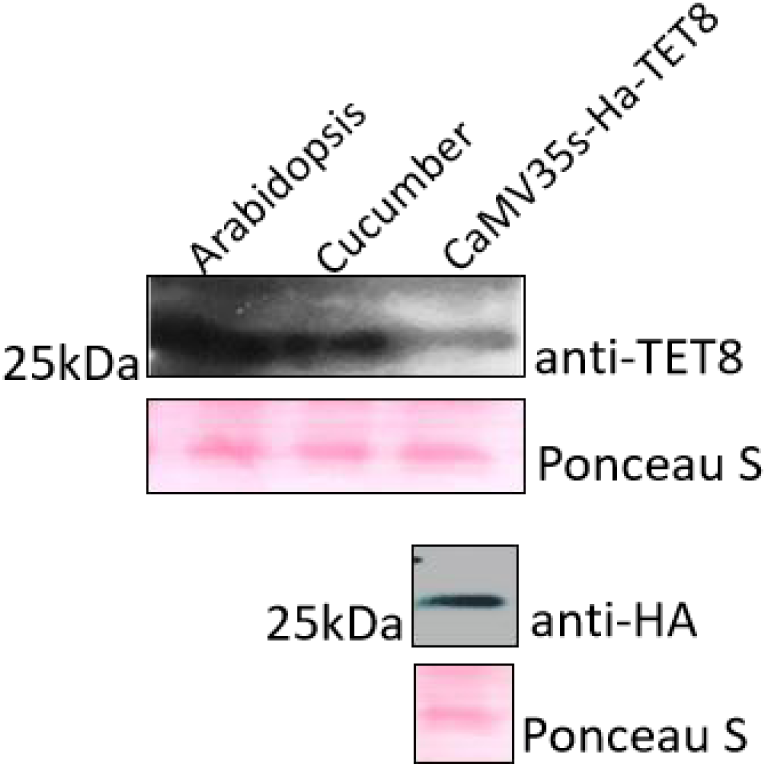
western blot of *N. benthamiana* cell lysate (CL) showing transient expression of HA-CsTET8 and arabidopsis and cucumber cell lysate as control showing 30kDa band corresponding to the size of TET8 with anti-TET8 antibody and anti-HA antibody.

**Fig. S16.**
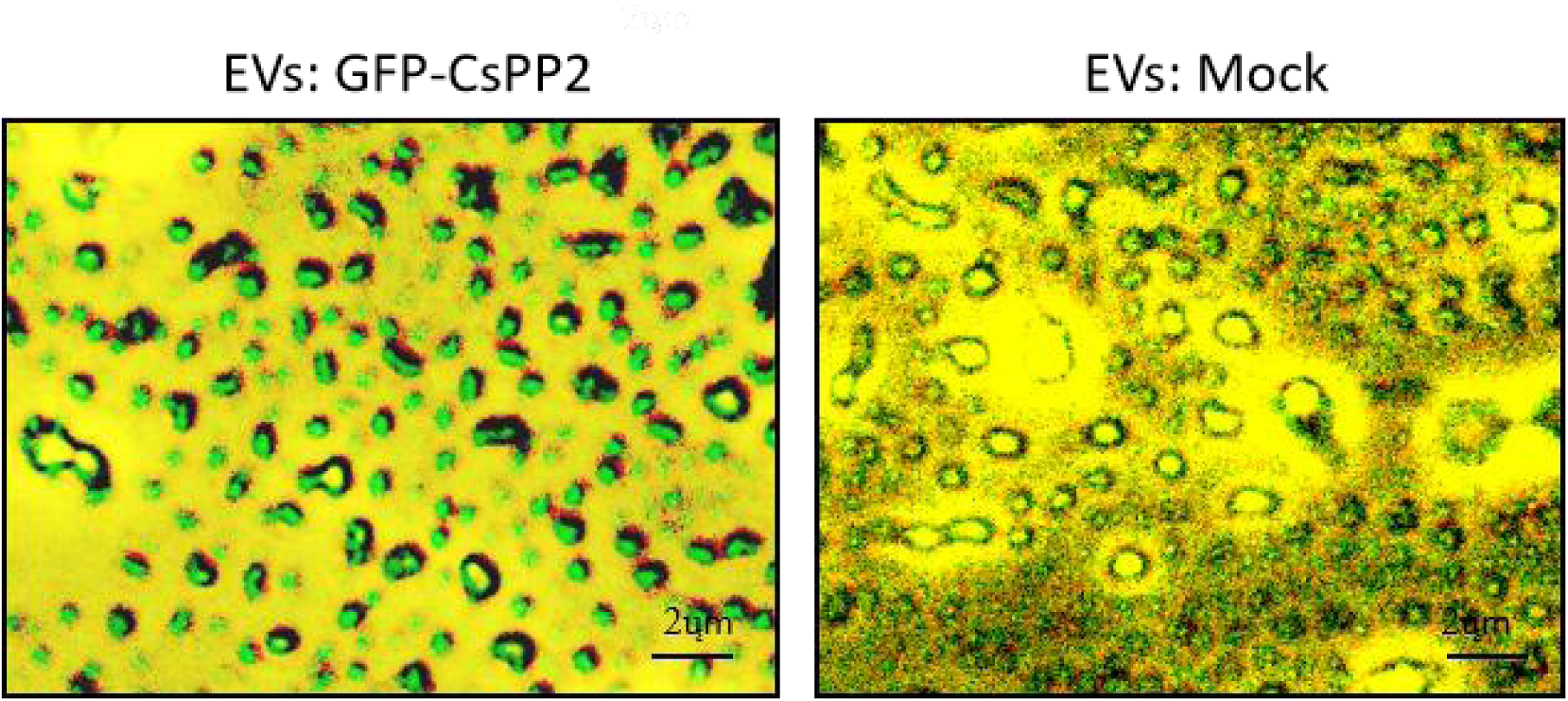
Image showing EVs from GFP-CsPP2 expressing *N. benthamiana* and mock plants as visualized under fluorescence microscope using GFP Filter The subcellular localization of CsTET8 was monitored by transiently expressing the CsTET8-pSITE2CA in *N. benthamiana* which revealed the cytoplasmic and plasma membranous localization of CsTET8.

**Fig. S17:**
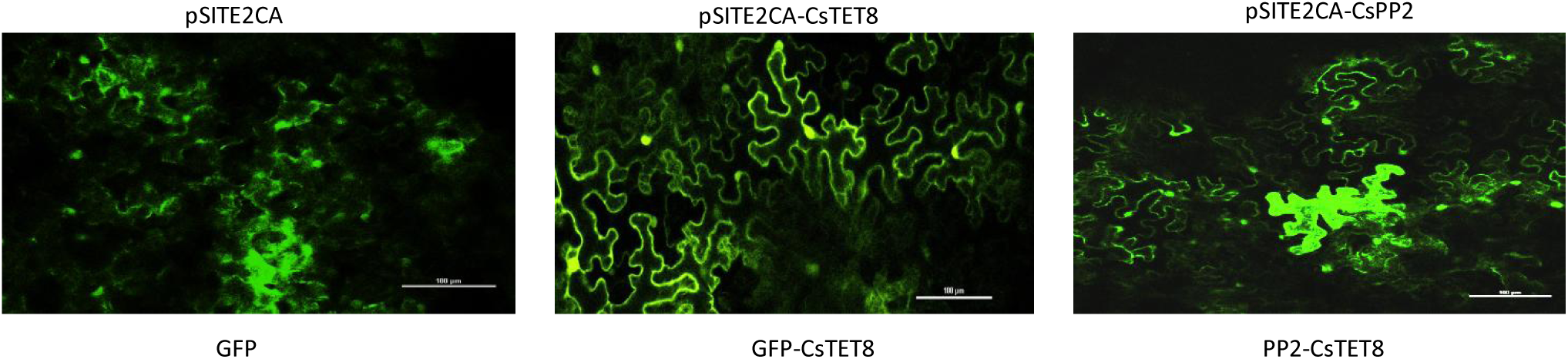
Localization of GFP-CsTET8 and GFP-CsPP2 in *N. benthamiana*. Speckles are observed on plasma membrane for both CsTET8 and CsPP2 hinting towards its association with membranous bodies.

**Fig. S18:**
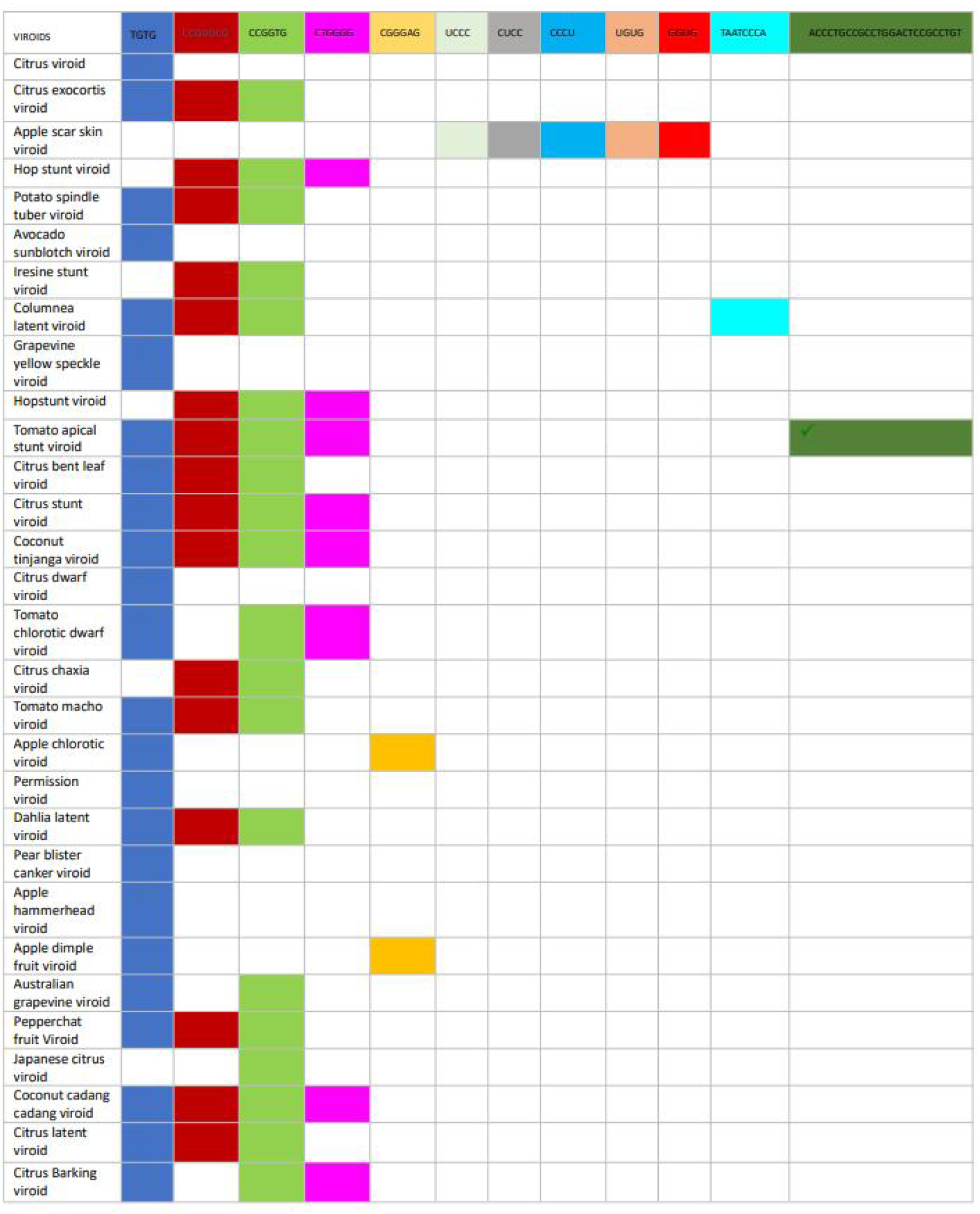
EXOmotifs observed in various viroids

**Fig. S19.**
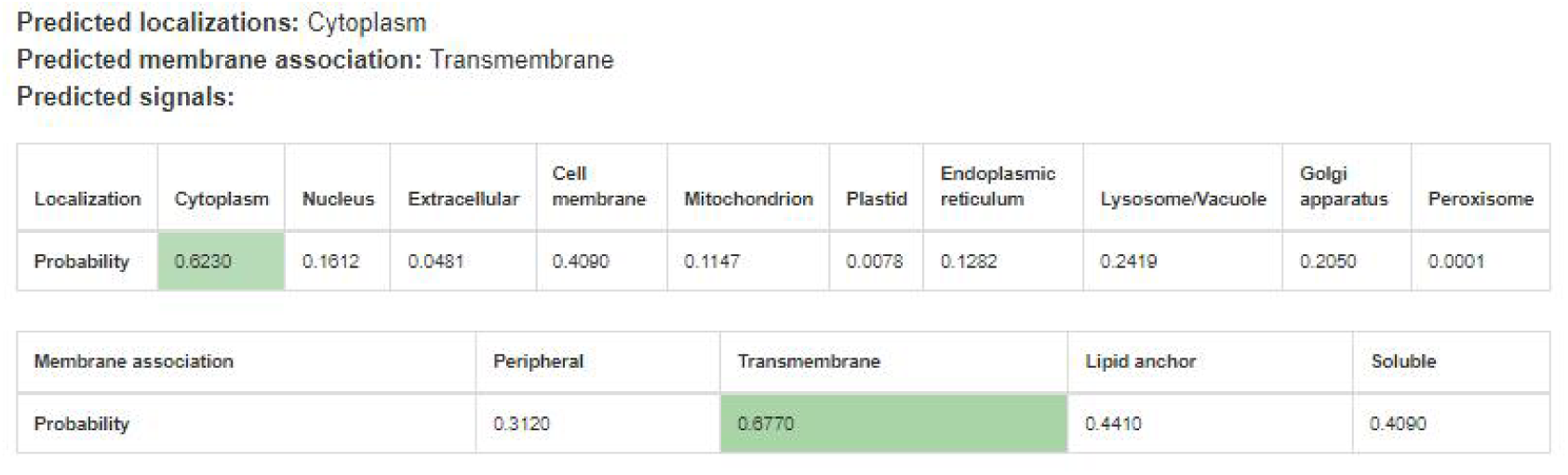
Showing the presence of transmemberane domain in the CsPP2 as analyzed using DeepLoC 2.1 tool (<u>DeepLoc 2.1 - DTU Health Tech - Bioinformatic Services</u>)

---

**Table S1:**
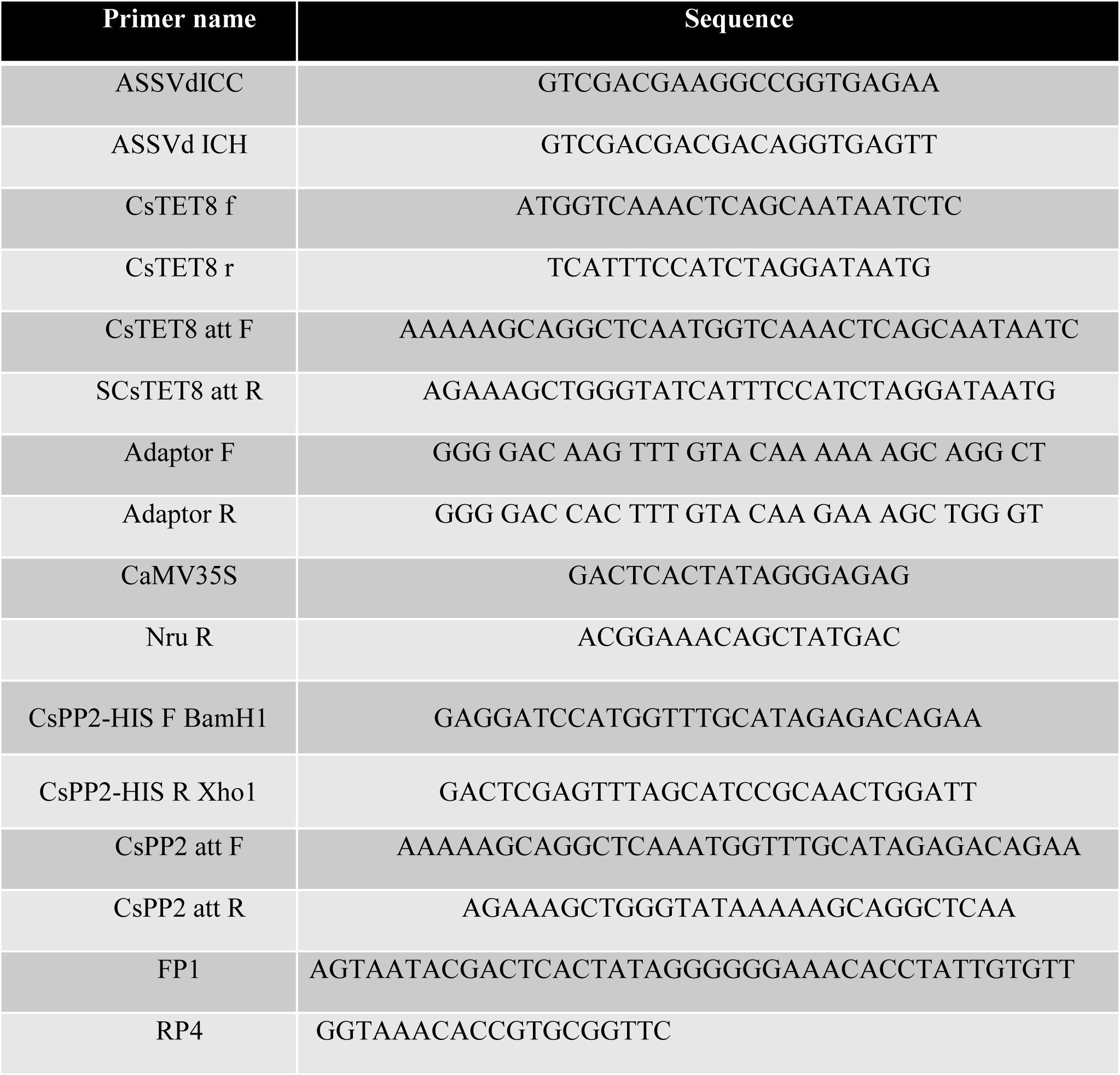
Primers used during the study.

